# Cell-to-cell signalling mediated via CO_2_: activity dependent axonal CO_2_ production opens Cx32 in the Schwann cell paranode

**DOI:** 10.1101/2025.04.04.647196

**Authors:** Jack Butler, Lowell Mott, Amol Bhandare, Angus Brown, Nicholas Dale

## Abstract

Loss of function mutations of Cx32, which is expressed in Schwann cells, cause X-linked Charcot Marie Tooth disease, a slowly progressive peripheral neuropathy. Cx32 is thus essential for the maintenance of myelin. During action potential propagation, Cx32 hemichannels in the Schwann cell paranode are thought to open and release ATP. As Cx32 hemichannels are directly sensitive to CO_2_, we have tested whether CO_2_ produced in the axon, as a consequence of the energetic demands of action potential propagation, might gate Cx32 hemichannels. Using isolated sciatic nerve from the mouse, we have shown that the critical components required for intercellular CO_2_ signalling are present (nodal mitochondria, the source of CO_2_; a CO_2_-permeable aquaporin, AQP1; paranodal Cx32; and carbonic anhydrase). We have used a membrane impermeant fluorescent dye FITC, which can permeate Cx32 hemichannels, to demonstrate the opening of Cx32 in Schwann cells in response to an external CO_2_ stimulus or during action potential propagation in the isolated nerve. Pharmacological blockade of AQP1 or allosteric enhancement of carbonic anhydrase activity greatly reduced Cx32 gating during action potential firing. By contrast, inhibition of carbonic anhydrase with acetazolamide greatly increased Cx32 gating. Cx32 gating was unaffected by the G-protein blocker GDPβS, indicating that it was not mediated by G protein coupled receptors. By expressing a modified Cx32 subunit, Cx32^DN^, that coassembles with Cx32^WT^, we have shown that the activity dependent dye loading of Schwann cells depends upon CO_2_ binding to Cx32. This CO_2_-dependent opening of Cx32 also mediates an activity dependent Ca^2+^ influx into the paranode and, by increasing the leak current across the myelin sheath, slows the conduction velocity. Our data demonstrate that CO_2_ can act via connexins to mediate neuron-to-glia signalling and that CO_2_ permeable aquaporins and carbonic anhydrase are key components of this signalling mechanism.

## Introduction

Connexin32 (Cx32) is expressed in Schwann cells and oligodendrocytes, the myelinating cells of the peripheral and central nervous system, respectively. These cells wrap around axons to form the myelin sheath, an insulating barrier that restricts voltage dependent ion fluxes to the nodes of Ranvier, thus enabling saltatory conduction (Huxley and Stampfli, 1949). Charcot Marie Tooth (CMT) disease is a slow progressing peripheral neuropathy that involves a loss of peripheral myelin integrity (Murakami et al., 1996). Typical symptoms of CMT include slowing of peripheral conductance velocity, loss of feeling in the extremities, *pes cavus* and in some cases muscle wasting. Mutations in the *gjb1* gene, encoding Cx32, result In the X linked version of CMT (CMTX) (Bergoffen et al., 1993; Fairweather et al., 1994; Hattori et al., 2003; Record et al., 2023). *Cx32*-null mice reproduce CMTX phenotypes, indicating CMTX is caused by a loss of Cx32 function (Scherer et al., 1998). This phenotype can be rescued by selective re-expression of Cx32 in Schwann cells, highlighting how fundamental Cx32 is to the maintenance of myelin health (Scherer et al., 2005).

Cx32 is a β connexin and is closely related to Cx26 and Cx30. Hemichannels of these connexin isoforms can be opened by increases in PCO_2_, at constant extracellular pH and physiological concentrations of Ca^2+^ (Huckstepp et al., 2010; Meigh et al., 2013; Dospinescu et al., 2019; Butler and Dale, 2023). CO_2_ sensitivity of these β connexins is dependent on a “carbamylation motif” (Meigh et al., 2013; Brotherton et al., 2022; Brotherton et al., 2024; Nijjar et al., 2025). While the mechanism of CO_2_ sensitivity has been most thoroughly studied in Cx26, Cx32 possesses the same carbamylation motif that is required for CO_2_ sensitivity (Dospinescu et al., 2019; Butler and Dale, 2023). In Cx32, CO_2_ carbamylates the primary amine of Lys124 in the motif. As this carbamylated amine is now negatively charged, it can form a salt bridge with Lys104 of the neighbouring subunit. The resulting carbamate bridges are thought to bias the hemichannel to the open state. Like most connexins, Cx32 will form gap junction channels where two hexamers of Cx32 in opposing membranes can dock together. Cx32 gap junction channels, however, are insensitive to the changes in PCO_2_ that can open Cx32 hemichannels (Dospinescu et al., 2019).

Myelin expresses Cx32 as both unopposed hemichannels in the paranodal membrane and also as reflexive gap junctions in the Schmidt Lanterman incisures (Bergoffen et al., 1993; Meier et al., 2004; Bortolozzi, 2018). The reflexive gap junctions provide radial diffusion pathways through the layers of myelin. However, radial diffusion pathways still exist in *Cx32*-null mice, suggesting a mechanism of redundancy or compensation (Balice-Gordon et al., 1998). Nevertheless, as *Cx32*-null mice still reproduce CMTX (Scherer et al., 1998), the loss of Cx32 hemichannel function must be sufficient to induce CMTX pathology.

Cx32 hemichannels in the paranode are thought to gate open and release ATP during action potential propagation (Nualart-Marti et al., 2013). The mechanism underlying the opening of Cx32 hemichannels in the paranode in response to action potential propagation remains uncertain. Two hypotheses have been proposed: i) because Cx32 is intrinsically voltage sensitive, hemichannel opening could be caused by transmembrane potential excursions during action potential propagation (Abrams et al., 2002); and ii) a rise in intracellular Ca^2+^ within the Schwann cell paranode possibly downstream of activation of a G-protein coupled receptor could open Cx32 (Carrer et al., 2018).

In this paper we explore an alternative hypothesis: that CO_2_, produced in the axon as a consequence of the energetic demands of restoring transmembrane ionic gradients following action potential propagation (via Na^+^/K^+^ ATPases), diffuses into the paranode to open Cx32. We have tested our hypothesis by careful consideration of the requirements of a CO_2_-based signalling system: a means of production (mitochondria); a channel to allow CO_2_ produced in the node to diffuse into the paranode as CO_2_ does not readily cross biological membranes; and a mechanism to terminate the actions of CO_2_ (carbonic anhydrase). We show that all of these components are present at the node/paranode and that their manipulation will alter the gating of Cx32 in ways that support our hypothesis.

## Results

### The components for CO_2_ signalling mediated via Cx32 are present in myelin

A CO_2_-based signalling system in myelin requires: a means of production; a channel to allow CO_2_ to cross the nodal and paranodal membranes; and a mechanism to terminate the actions of CO_2_ (Fig 1). It is already known that: mitochondria are present in the node (Ohno et al., 2011); Cx32 is expressed in the paranode (Bergoffen et al., 1993); AQP1, highly permeable to CO_2_ (Endeward et al., 2006; Musa-Aziz et al., 2009), is expressed in Schwann cells (Gao et al., 2006; Segura-Anaya et al., 2015); and carbonic anhydrase is universally present in every cell. Here we have used high resolution microscopy to examine the precise subcellular localisation of these components relative to each other alongside markers of the nodal and paranodal regions (Fig 2 figure supplement 1).

**Figure 1.**
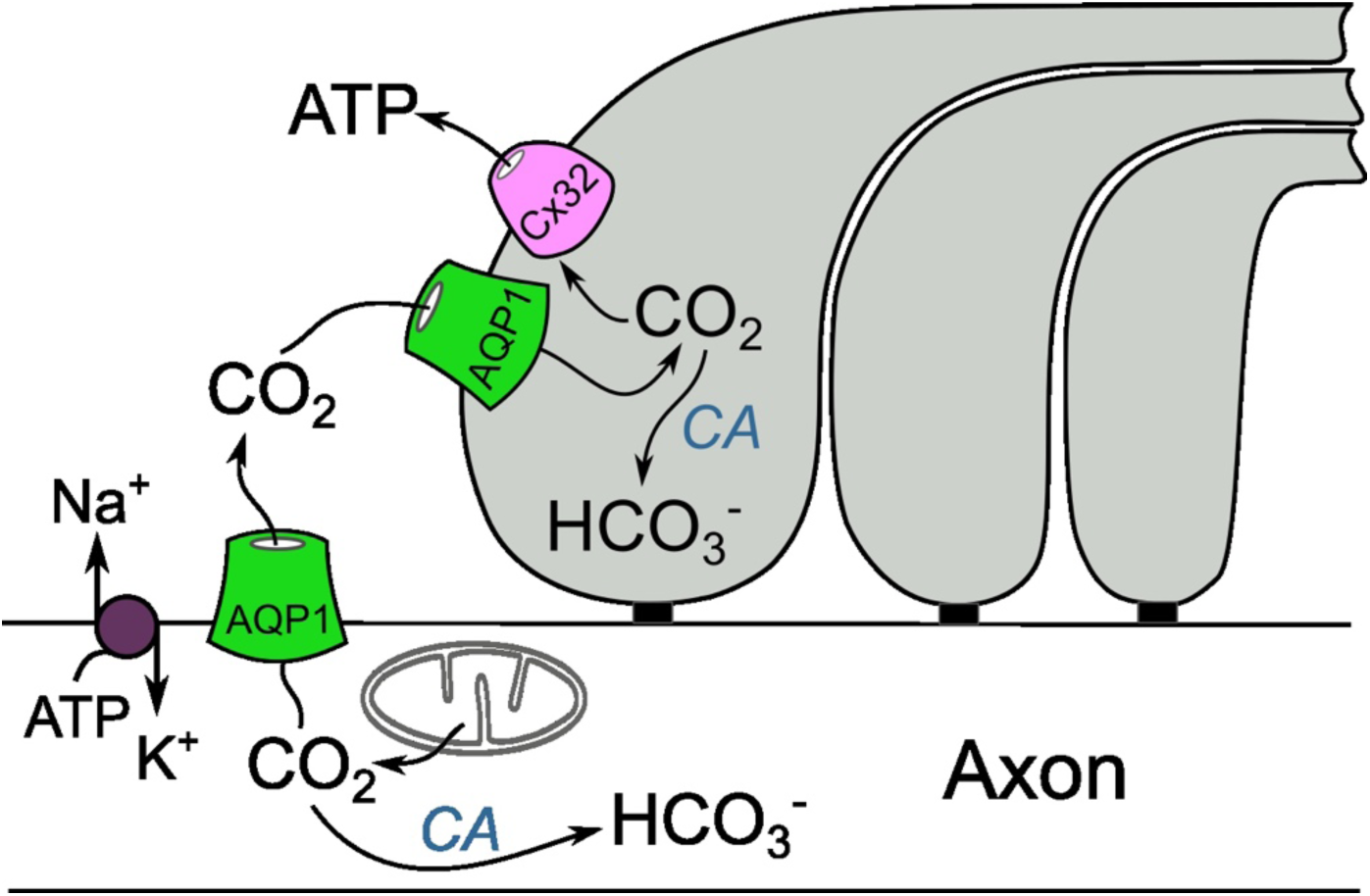
Hypothesised Cx32 mediated CO_2_ signalling cascade in peripheral myelin. Three fingers of a myelin paranode have been used for illustrative purposes. Restoration of transmembrane ionic gradients following action potential propagation via the actions of Na^+^/K^+^ ATPases incurs a metabolic cost and increases production of ATP and CO_2_. AQP1, permeable to CO_2_, provides a pathway for CO_2_ to leave the node and enter the paranode and bind to Cx32 on the intracellular loop. This triggers opening of Cx32 and release ATP. Carbonic anhydrase(CA) catalyzes the combination of CO_2_ and H_2_O and ultimately the production of HCO_3_^-^ and H^+^ ions and effectively competes with Cx32 for CO_2_.

**Figure 2.**
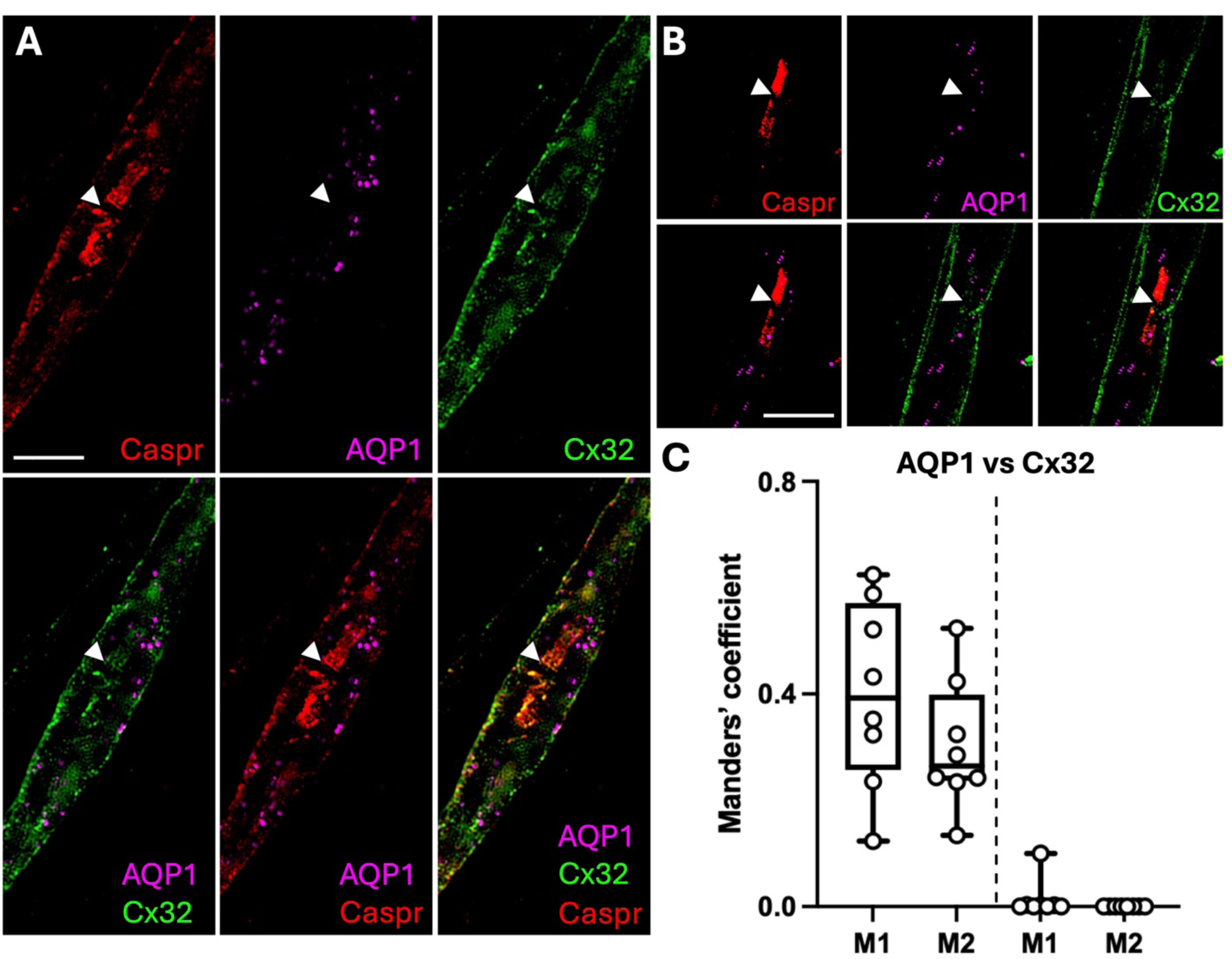
AQP1 localizes to both the Schwann cell paranode and also the axonal node. **A,** B) Representative confocal SIM images from a single optical plane showing the localisation of Caspr, Cx32 and AQP1 in isolated mouse sciatic nerve. Arrowheads indicate the node. Scale bars, 10 μm. C) Boxplots showing degree of colocalization between Cx32 and AQP1 at the node/paranode. Control measurements (to right of dashed line) used these same images with one channel flipped 90° and the same thresholds as when measuring colocalization. Kruskal Wallis ANOVA p < 0.0001.

#### AQP1 is present in the axonal node and Schwann cell paranode

AQP1, a CO_2_ permeable aquaporin (Endeward et al., 2006; Musa-Aziz et al., 2009), was localised to the Schwann cell paranode and outer myelin membrane (Fig 2). APQ1 expression also colocalised with Caspr, showing that it was present in the axonal nodal membrane (Einheber et al., 1997). Interestingly, analysis of colocalization Cx32 and AQP1 in the node/paranode region showed that AQP1 was in close proximity to Cx32 in the paranode (M1: mean 0.400; 95% CI 0.254 to 0.546 and M2: mean 0.301; 95% CI 0.199 to 0.403). This subcellular localisation of AQP1, would allow it to act as a conduit for CO_2_ generated at the axonal node to enter into the Schwann cell paranode and interact with Cx32.

#### Mitochondria localise to the axonal node and Schwann cell paranode and may be brought in close proximity to Cx32 via SFXN1

SFXN1 is a mitochondrial protein that also binds to Cx32 (Fowler et al., 2013). Using cytochrome C (CytC), as a mitochondrial marker we found that mitochondria were localised in both the axonal node and Schwann cell paranode (Fig 2 figure supplement 2), in accordance with previous reports (Rydmark et al., 1998; Ohno et al., 2011). There was colocalization between Cx32 and CytC in the Schwann cell paranode (Fig 2 figure supplement 2, mean; 95% confidence interval, M1: 0.314; 0.198 to 0.431 and M2: 0.261; 0.165 to 0.357). There was also colocalization in the Schwann cell paranode between CytC and SFXN1 (Fig 2 figure supplement 2, M1: 0.568; 0.441 to 0.695 and M2: 0.462; 0.336 to 0.588). This suggests that SFXN1 may facilitate the association of Cx32 and mitochondria (Fowler et al., 2013). AQP1 also closely associated with CytC (Fig 2 figure supplement 3). Interestingly, SFXN1 was also observed in the absence of CytC (Fowler et al., 2013) suggesting that it has additional cellular roles unrelated to its mitochondrial function.

#### Carbonic anhydrase is present in the paranode

We observed strong expression of CAII in non-myelinated fibres (Fig 3). However, consistent with earlier reports (Cammer and Tansey, 1987) we also observed weaker but more localised expression in myelinated fibres, specifically at the axonal node and Schwann cell paranode (Fig 3).

**Figure 3:**
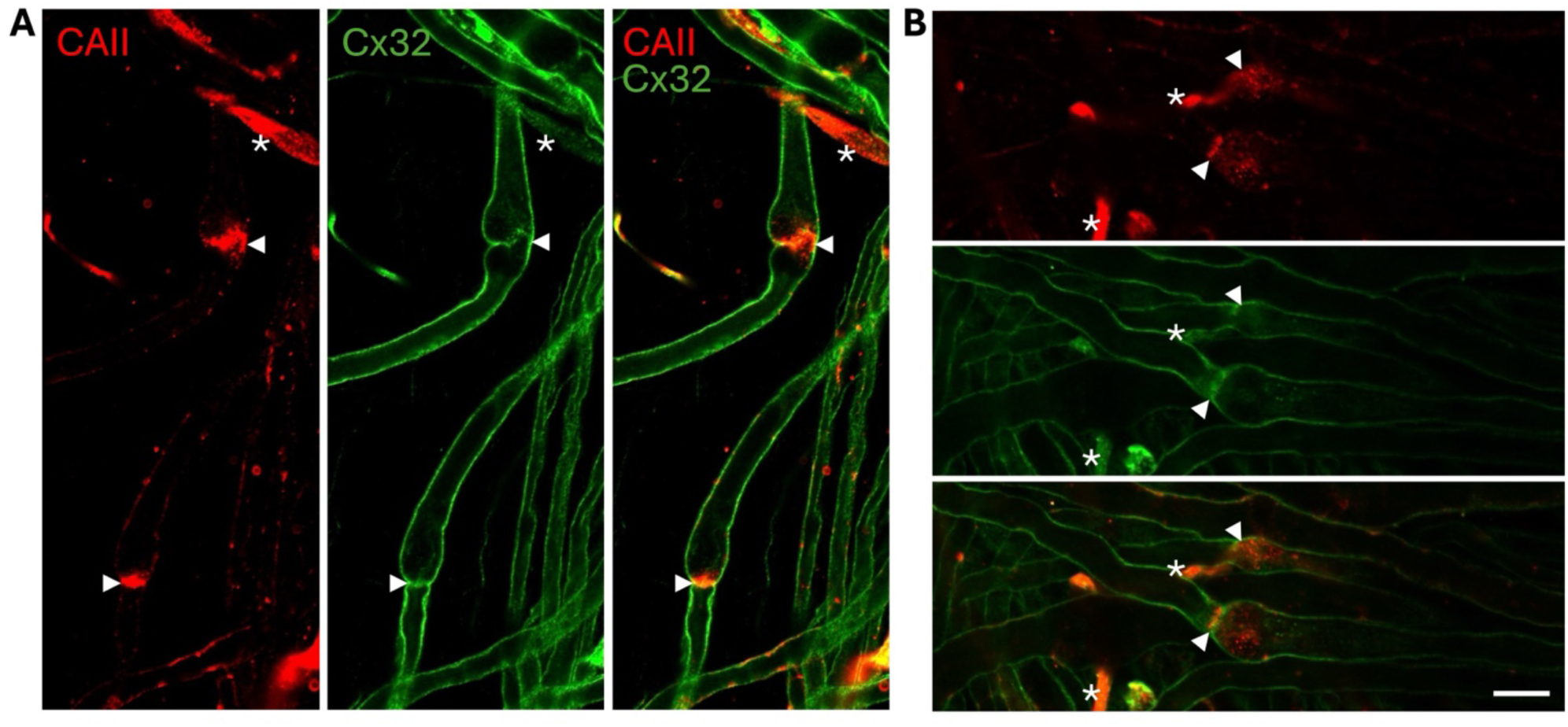
CAII localizes to myelinating Schwann cells, in particular to the axonal node and the Schwann cell paranode. (A and B) Representative confocal LSM images in single optical plane showing the localisation of CAII and Cx32 in isolated mouse sciatic nerve. Arrow heads indicate the node. Intense CAII staining, denoted by a white asterisk (*) is present in non-myelinated fibres. Scale bar applies to A and B: 10μm.

### CO_2_-dependent dye loading of Schwann cells in sciatic nerve

We first examined whether Cx32 hemichannels in Schwann cells could be opened by application of hypercapnic aCSF. We exposed isolated sciatic nerves to FITC in aCSF at different levels of PCO_2_. As FITC is membrane impermeant but can readily move through channels with large pores such as Cx32 hemichannels (Butler and Dale, 2023), any CO_2_-dependent dye loading would thus indicate gating of a CO_2_-sensitive large pore channel.

At 35 mmHg, a level of PCO_2_ that is too low to open Cx32 hemichannels (Huckstepp et al., 2010; Dospinescu et al., 2019), FITC loading was not observed (Fig 4). However, in the presence of hypercapnic aCSF (70mmHg, sufficient to open Cx32 hemichannels) dye loading into the paranode and outer myelin layers was readily observed (Fig 4, p < 0.0001, compared to 35 mmHg). Note that the axons did not load with FITC.

**Figure 4:**
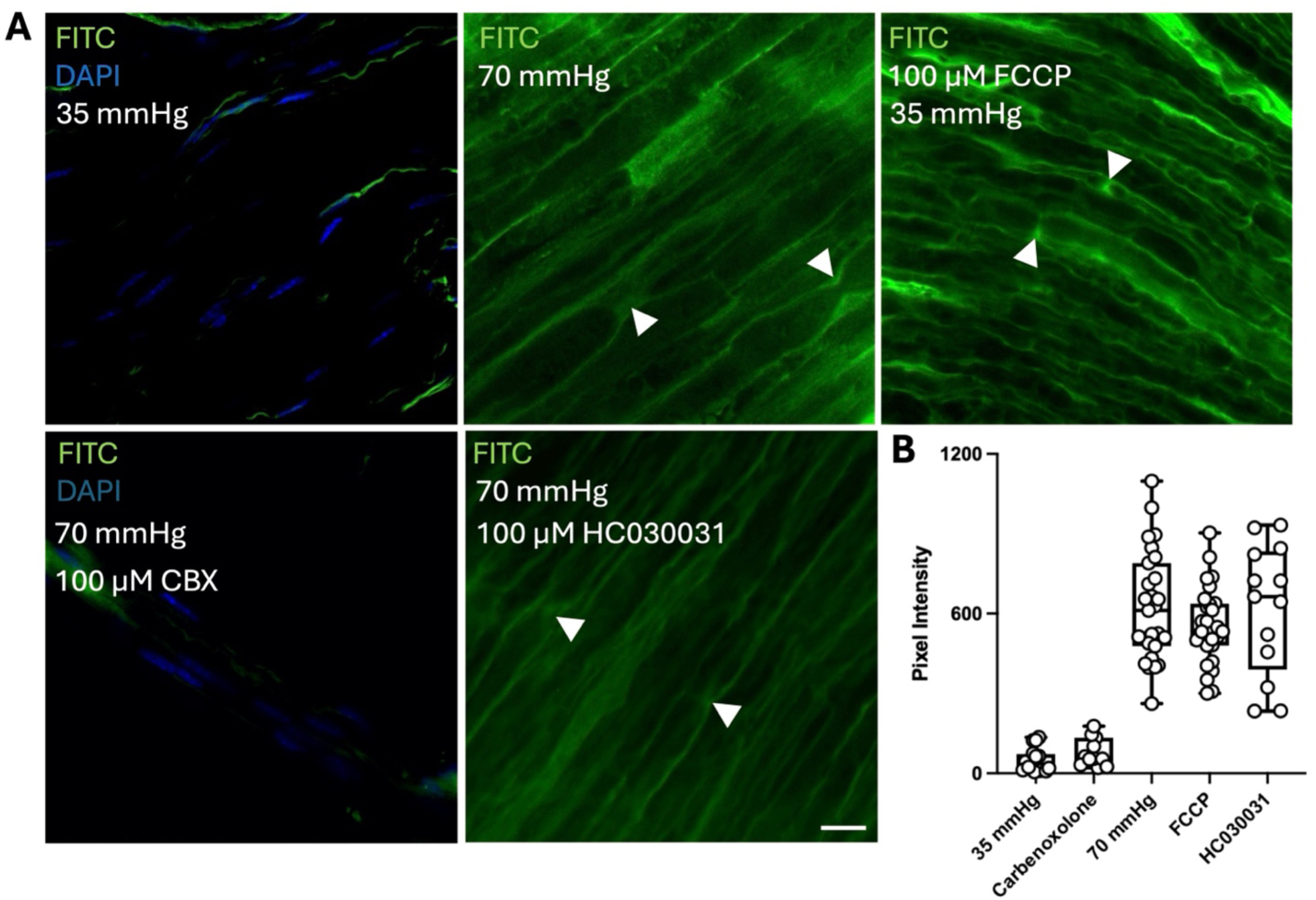
A membrane impermeable dye, FITC, loads into Schwann cell paranodes in a CO_2_ dependent manner through a hemichannel. A) Representative images showing the FITC loading into mouse sciatic nerve bundles. Arrow heads indicate the node. Little FITC loading occurs in response to control (35mmHg, 10 mins) aCSF. FITC loading was greatly increased by 70 mmHg aCSF (10 mins) and application of 100 μM FCCP. FITC loading in 70 mmHg aCSF was blocked by carbenoxolone (CBX) but not the TRPA1 antagonist HC030031. B) Boxplot showing intensity of FITC fluorescence under the different conditions. Each point represents a separate ROI from 5 different nerves for each condition. Scale bar – 10μm.

To demonstrate that mitochondrially produced CO_2_ could gate Cx32, we used a mitochondrial uncoupler, FCCP, to maximise rates of endogenous CO_2_ generation (Balboni and Lehninger, 1986). We found that FCCP, applied in 35 mmHg aCSF, caused significantly increased dye loading into Schwann cell paranodes and outer myelin layers (p < 0.0001) compared to nerves loaded at 35 mmHg PCO_2_ with no FCCP (Fig 4).

CO_2_-evoked FITC loading was abolished in the presence of carbenoxolone, indicating FITC entry occurred through a carbenoxolone sensitive hemichannel (p < 0.0001, Fig 4). TRPA1 can open with intracellular acidification (Wang et al., 2010), however CO_2_-evoked FITC loading was not blocked by a specific TRPA1 antagonist, HC030031, supporting that dye entry occurred via a connexin rather than TRPA1 (p = 0.8643, Fig 4).

Schwann cells express two connexins: Cx32 and Cx31.3 (also known as Cx29) (Jeng et al., 2006; Gerber et al., 2021). Cx31.3 lacks the carbamylation motif and is therefore unlikely to be CO_2_ sensitive. To confirm this, we measured ATP release via Cx31.3 expressed in HeLa cells (Liang et al., 2011) in response to changes in PCO_2_ and membrane depolarisation, by means of a co-expressed genetically encoded sensor, GRAB_ATP_. HeLa cells transfected only with GRAB_ATP_ but not Cx31.3 did not show any fluorescent changes in response to 70 mmHg PCO_2_ or 50 mM K^+^ (Fig 4 fig supplement 1). However, in cells transfected with Cx31.3, 50 mM KCl induced ATP release (Fig 4 fig supplement 1). By contrast, a stimulus of 70 mmHg PCO_2_ was ineffective at triggering ATP release (Fig 4 fig supplement 1). We have previously shown that this level of PCO_2_ readily induces ATP release via Cx32 (Butler and Dale, 2023; Lovatt et al., 2025). This confirms that Cx31.3 is not sensitive to CO_2_ and makes it most likely that the CO_2_-dependent entry of FITC into the Schwann cells was via Cx32.

### Activity-dependent loading of FITC into Schwann cells depends on CO_2_ production

To test whether Cx32 might open and permit FITC entry into the paranode during action potential propagation, we bathed isolated nerves in aCSF (35 mmHg) and stimulated them electrically at 30 Hz, while measuring the compound action potential (CAP). Upon electrical stimulation, FITC entry into myelin was observed (Fig 5). We confirmed that FITC loaded into paranodes by counterstaining with the paranode marker Caspr (Einheber et al., 1997) (Fig 5). FITC did not load into the axons. FITC loading into myelin was correlated positively with the stimulus duration (Fig 6B). Electrical stimulation of the axon was required for dye loading as it did not occur in nerves that were exposed to FITC for 10 mins in the absence of stimulation (Fig 4 A,B).

**Figure 5:**
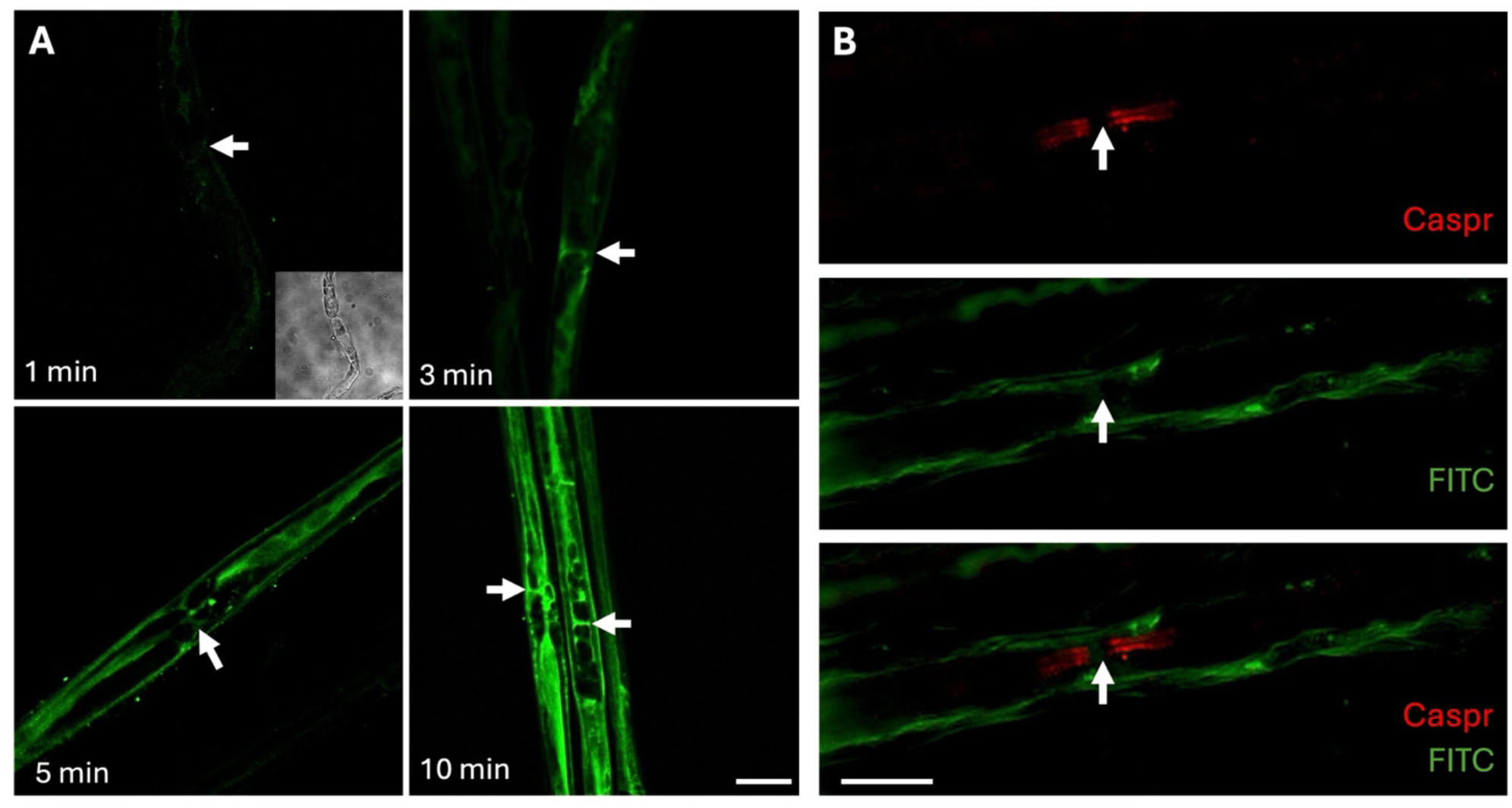
Activity dependent loading of FITC into Schwann cell paranodes. A) Representative images showing the FITC loading into mouse sciatic nerve bundles in response to different lengths of stimulation (30 Hz). Arrows indicate the paranode. B) An isolated mouse sciatic nerve fibre loaded with the membrane impermeable dye FITC (30 Hz, 5 min) and counterstained with Caspr, an axonal membrane protein which is expressed only in the paranodal region. White arrows indicate the paranode of interest. Note lack of loading into the axon. Scale bars – 15 μm.

**Figure 6:**
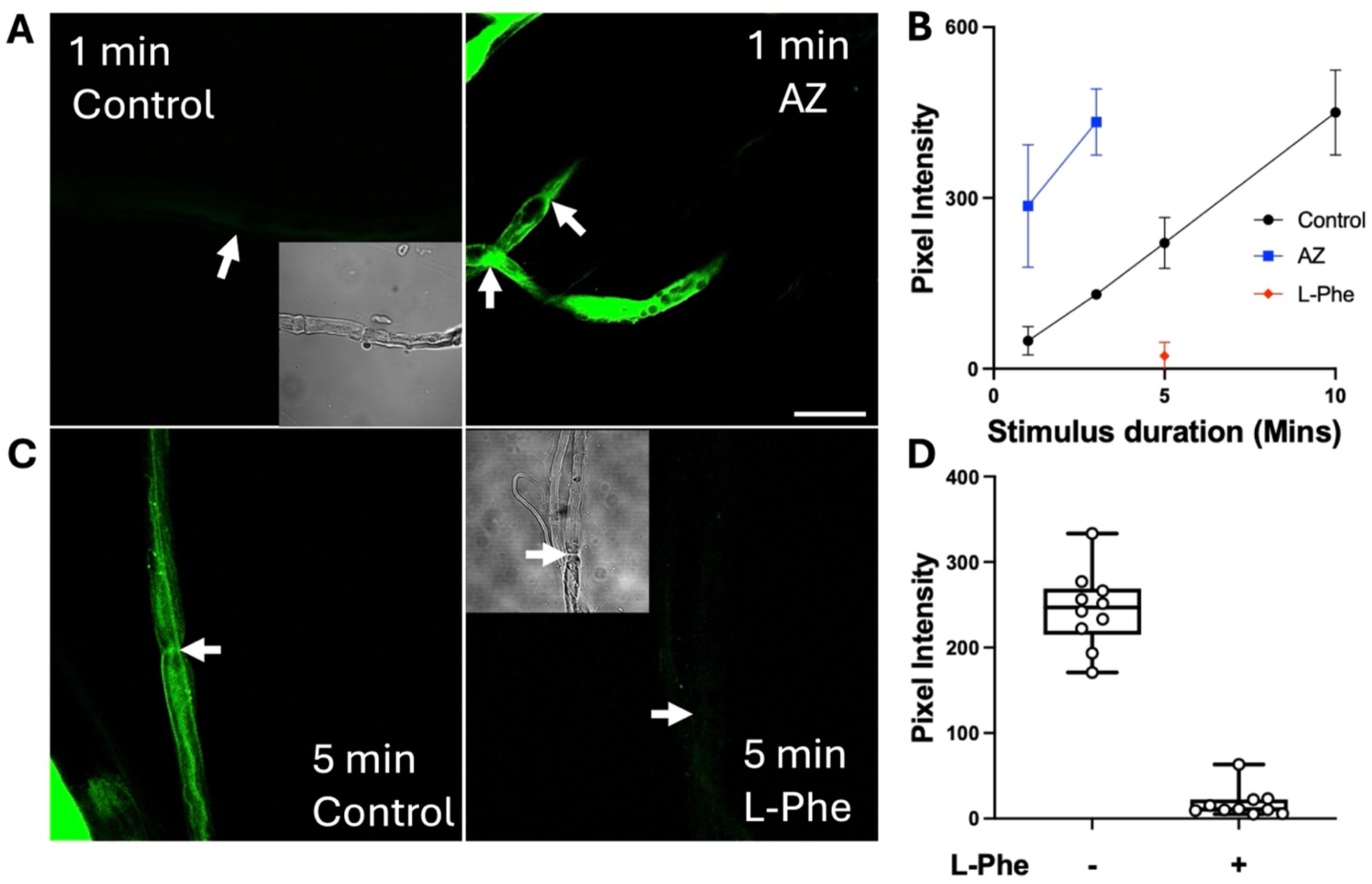
Activity dependent loading of FITC is sensitive to manipulation of carbonic anhydrase activity. A) Representative images showing the FITC loading into mouse sciatic nerve bundles in response to 1 min of electrical stimulation in the absence (left) or presence (right) of the carbonic anhydrase inhibitor, acetazolamide. Arrows indicate the position of paranodes. Brightfield inset shows the presence of the nerve fibre and the arrowhead in the fluorescence image indicates its position. B) Summary plot showing how the pixel intensity of Schwann cell paranodes, and therefore FITC loading, vary in response to stimulus duration and the presence of acetazolamide (each point mean ± SD). C) Representative images showing the FITC loading into mouse sciatic nerve bundles in response to five minutes of stimulation in the absence (left) or presence (right) of the carbonic anhydrase enhancer, L-Phenylalanine. Brightfield inset shows the presence of the nerve fibre and the arrowhead in the fluorescence image indicates its position. D) Boxplot showing the effect L-Phenylalanine had on FITC loading into mouse Schwann cell paranodes. Scale bars – 15 µm. Each point represents a separate ROI from 5 different nerves. L-Phe vs control MW test: p<0.0001.

To test whether this activity dependent FITC loading was also CO_2_ dependent, we first manipulated the activity of carbonic anhydrase (CA). Inhibition of CA activity, via acetazolamide, should increase the local PCO_2_ as the conversion of CO_2_ to carbonic acid will be slowed. We found that acetazolamide (100 µM) greatly increased FITC loading into the Schwann cell paranode in response 30 Hz stimulation for 1 or 3 minutes (p=0.001 and p=0.0121 respectively, Fig 6A,B).

L-phenylalanine (L-Phe) is an allosteric enhancer of CA activity (Temperini et al., 2006). Myelinating Schwann cells express SLC7A5 (Gerber et al., 2021; Karlsson et al., 2021), the gene that encodes the L-type amino acid transporter. As this transports L-Phe (Nguyen et al., 2021), bath application of L-Phe (1 mM) to isolated nerve, should be effective in enhancing the activity of intracellular CA in Schwann cells. The accelerated conversion of CO_2_ to carbonic acid in the presence of L-Phe would be expected to reduce activity dependent dye loading. We indeed observed that treatment with L-Phe greatly reduced activity dependent FITC loading into the Schwann cell paranode (p = 0.0159, Fig 6C,D). Neither acetazolamide nor L-Phe altered the amplitude or the current-amplitude curves of the CAP (Fig 6, fig supplement 1) indicating that these drugs did not affect the excitability of the axon.

CO_2_ is produced from the Krebs cycle during two steps of oxidative decarboxylation. The first step of the Krebs cycle requires isomerisation of citrate to isocitrate, via the enzyme aconitase, to enable the first decarboxylation event (via isocitrate dehydrogenase). In neurons aconitase can be selectively blocked by 50 µM H_2_O_2_ (Tretter and Adam-Vizi, 2000). We therefore tested whether this dose of H_2_O_2_ could reduce activity dependent FITC loading into the Schwann cell. Application of H_2_O_2_ greatly reduced FITC loading (Fig 7). There was no effect of H_2_O_2_ on the CAP (Fig 7, figure supplement 1), supporting the notion that Cx32 gating depends upon the Krebs cycle and production of CO_2_.

**Figure 7.**
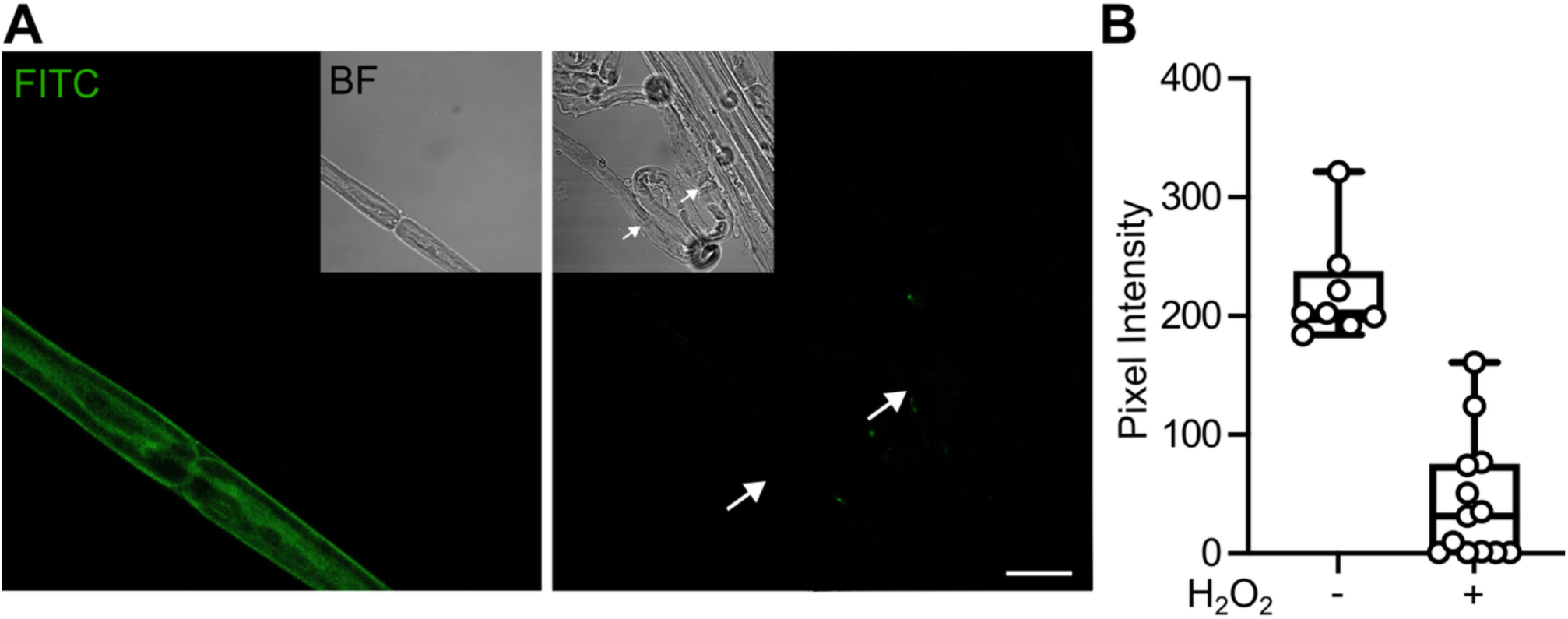
Activity dependent loading of the FITC is reduced by inhibition of the Krebs cycle. A) Representative images showing the FITC loading into mouse sciatic nerve bundles in response to 5 minutes of 30 Hz stimulation in the absence (left) or presence (right) of the 50 µM H_2_O_2_ which blocks aconitase and the Krebs cycle. Brightfield inset shows the presence of the nerve fibre and the arrows in the fluorescence images indicates position of paranodes. Scale bar – 15 μm. B) Boxplot showing the effect 50 µM H_2_O_2_ had on FITC loading into mouse Schwann cell paranodes. Each point represents a separate ROI from 5 different nerves. H_2_O_2_ vs control MW test: p<0.0001.

As a final test of our hypothesis that activity dependent CO_2_ production in the axon gates Cx32 in the paranode, we used a specific blocker of AQP1, TC AQP1-1 (80 µM, (Ghosh et al., 2020)). We found blockade of AQP1 greatly reduced FITC loading into the Schwann cell paranode following 5 minutes of stimulation at 30 Hz, compared to that of WT (p< 0.0001, Fig 8). This supports our hypothesis and also indicates that AQP1 is a key conduit for CO_2_ to diffuse from the axonal node to the Schwann cell paranode. TC APQ1-1 had no effect on the amplitude or the current-amplitude curves of the CAP (Fig 8, figure supplement 1) indicating that it did not affect either the excitability of the axon or its capacity for action potential generation.

**Figure 8.**
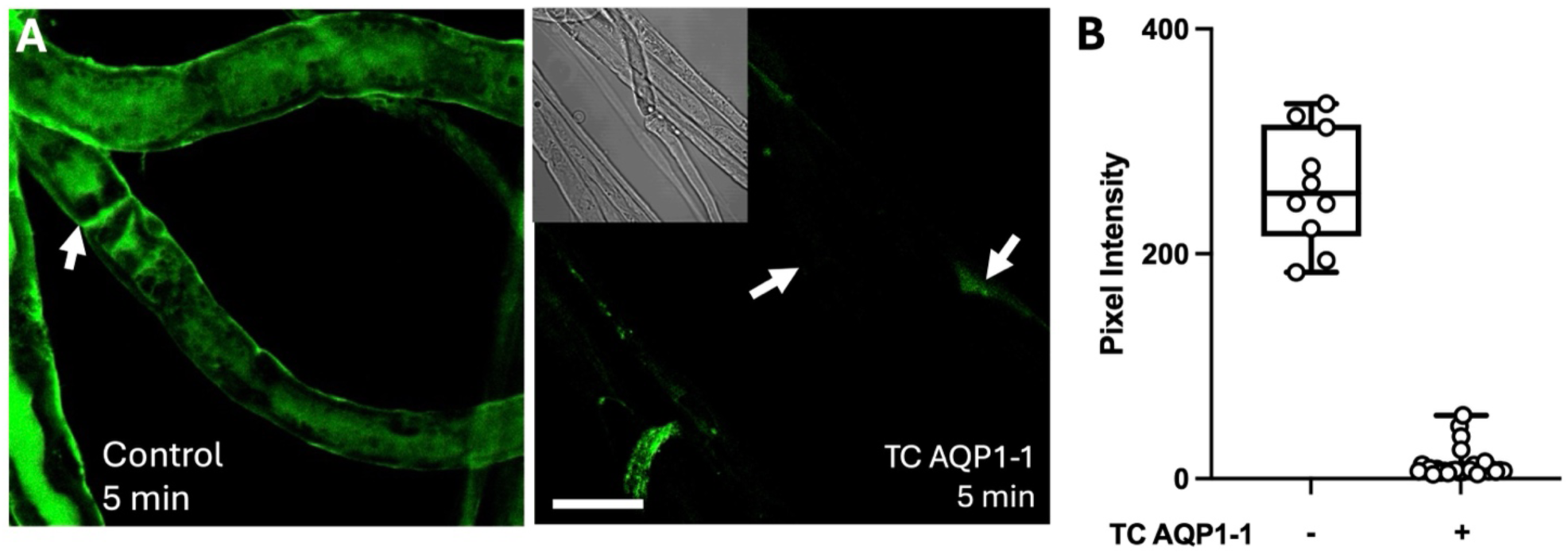
Activity dependent loading of the FITC is reduced by inhibition of AQP1. A) Representative images showing the FITC loading into mouse sciatic nerve bundles in response to 5 minutes of 30 Hz stimulation in the absence (left) or presence (right) of the AQP1 blocker TC AQP1-1. Brightfield inset shows the presence of the nerve fibre and the arrows in the fluorescence images indicates position of paranodes. Scale bar – 15 μm. B) Boxplot showing the effect TC AQP1-1 had on FITC loading into mouse Schwann cell paranodes. Each point represents a separate ROI from 5 different nerves. TC AQP1-1 vs control MW U test: p<0.0001.

While our data are consistent with activity dependent CO_2_ production in the node gating Cx32 in the paranode, they do not eliminate the possible involvement of other signalling pathways such as those mediated by G-protein coupled receptors (GPCRs). We therefore used GDPβS (100 µM) as a general blocker G-protein mediated signalling. As a positive control we showed that application of GDPβS blocked ATP receptor mediated increases in intracellular Ca^2+^ in the paranode (Fig 8, figure supplement 2). However, the application of GDPβS had no effect on activity dependent FITC loading (Fig 8, figure supplement 2).

#### Enhancement of FITC loading by block of CA is not mediated by pH changes

Inhibition of CA by acetazolamide could plausibly lead to subsequent alkalosis, as the production of HCO_3_^-^ and H^+^ ions will be reduced. We therefore tested whether alkalosis by itself was sufficient to enhance activity dependent FITC loading by applying NH_4_Cl (100 µM), but this had no effect (p = 0.1257, Fig 8 figure supplement 3).

We quantified the changes in intracellular pH induced upon perfusion of acetazolamide or NH_4_Cl by using the pH sensitive dye BCECF (Fig 8 figure supplement 4). We found that NH_4_Cl induced greater increases in intracellular pH (change (median; 95% CI): 0.1579; 0.119 to 0.1968), than did acetazolamide which had no significant effect on intracellular pH (change: - 0.0147; -0.040 to 0.011). The enhancement of activity dependent FITC loading by acetazolamide cannot therefore be explained by changes in intracellular pH.

#### Activity dependent loading of FITC depends on CO_2_ binding to Cx32

To directly address both the involvement of Cx32 and specifically binding of CO_2_ to Cx32 via the carbamylation motif, we utilised a dominant negative subunit, Cx32^DN^. Cx32^DN^ carries the K124R and K104A mutations and can thus neither bind CO_2_ nor form a salt bridge with a neighbouring subunit that has bound CO_2_. We have previously shown that Cx32^DN^ removes CO_2_ sensitivity from cells that express Cx32^WT^ (Butler and Dale, 2023). Using acceptor depletion FRET (Gu et al., 2004; van de Wiel et al., 2020) we documented that Cx32^DN^ coassembles with Cx32^WT^ (Fig 9, figure supplements 1 and 2). Furthermore, Cx32^DN^ formed gap junctions that retained their permeability to small molecules (Fig 9 figure supplement 3).

**Figure 9.**
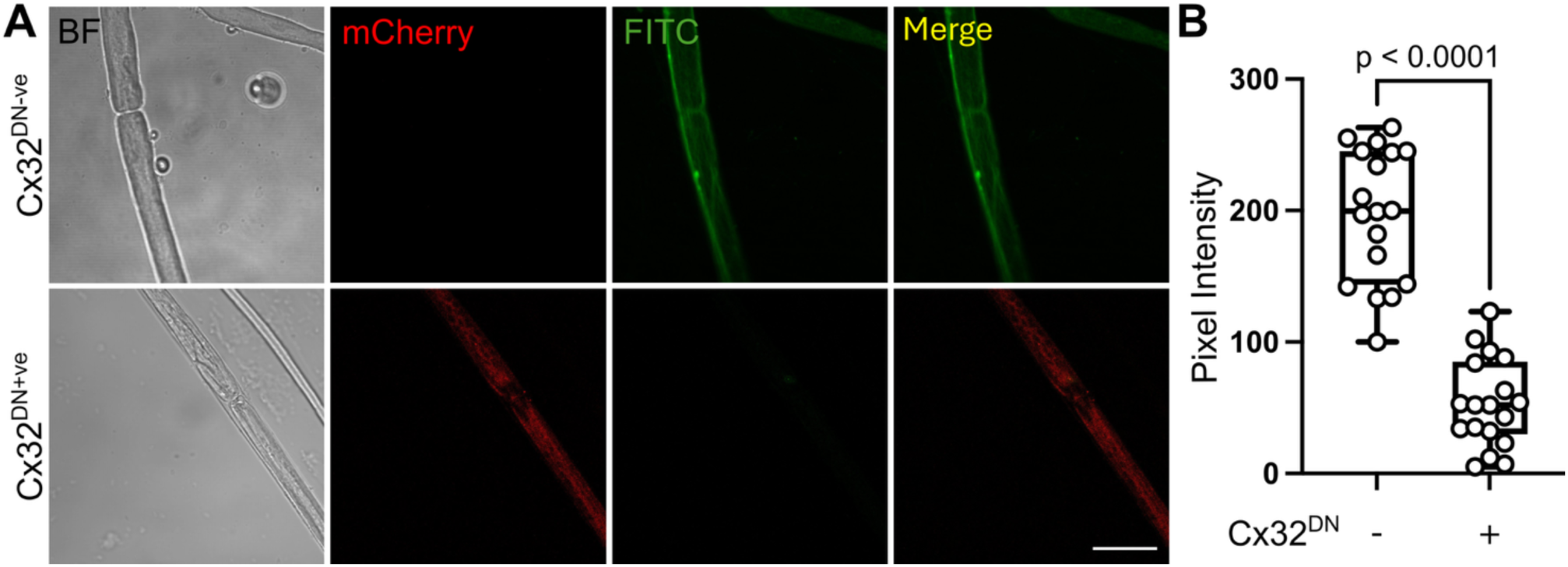
Activity dependent dye loading into Schwann cells depends on CO_2_ binding to Cx32. A) Images of single axons dissected from a sciatic nerve transduced with AAV-Mpz-Cx32^DN^-IRES-mCherry. Axons that do not express Cx32^DN^ do not exhibit mCherry fluorescence and show robust dye loading to 15 Hz stimulation. By contrast axons that express mCherry are Cx32^DN+ve^ and do not show dye loading during stimulation. Scale bar 15 µm. B) Summary graph showing the pixel intensity of axons that are Cx32^DN+ve^ versus the control Cx32^DN-ve^. MW p<0.0001, each circle an individual paranode from n=5 nerves.

We therefore transduced sciatic nerve with AAV-Mpz-Cx32^DN^-IRES-mCherry. This construct design uses the Mpz promoter to restrict expression to the Schwann cell. The Cx32^DN^ gene is not tagged at the C-terminus but, as there is an IRES-mCherry sequence, cytosolic expression of mCherry permits identification of the transduced Schwann cells (van de Wiel et al., 2020). Expression of Cx32^DN^ greatly reduced activity dependent dye loading into Schwann cells that expressed Cx32^DN^ but not in those that did not in the same nerve (Fig 9).

### A simplified model of the paranode supports CA as a key regulator of Cx32 gating

To gain further insight, we made a simplified model of the paranode (as a single cell that in effect incorporated the nodal mitochondrion) to explore the effects of CA activity on loading of FITC into Schwann cell paranodes (see Methods and Fig 10). The mitochondrion in this simplified “paranode” was based on a model proposed by Matsuda et al., (2020). The Matsuda model, which accurately replicates the experimentally observed dynamics of ATP production in mitochondria of myotubes, incorporates the concept of mitochondrial priming: that electrical activity in the myotube enhances the rate of ATP synthesis.

**Figure 10:**
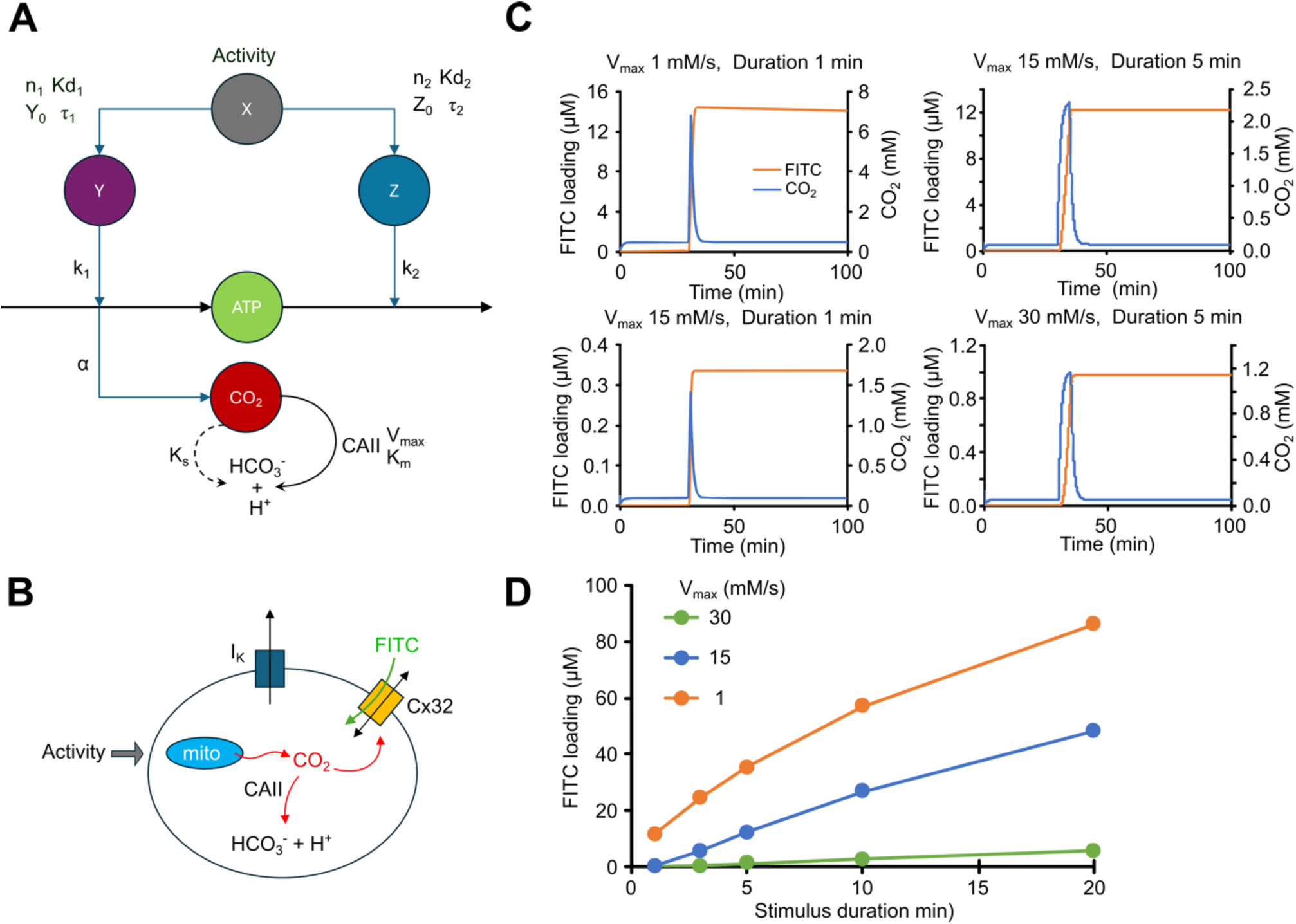
Simple model of CO_2_ signalling at the paranode reproduces experimentally observed patterns of activity dependent FITC loading. A) Adaptation of the Matsuda et al model to incorporate CO_2_ production and metabolism via CAII. Action potentials provide the input to the model as activity variable X, which determines variables Y and Z, which determine ATP production and consumption. CO_2_ is proportional to ATP production, via the rate constant α. Code for the model is provided in the Matlab files CO2Cx32Model.m and CO2Cx32Plot.m. B) Incorporation of the modified Matsuda et al model into a single cell that possesses a K^+^ leak channel and Cx32 and represents the paranode, albeit including the nodal mitochondria. (C) Outputs from the model to show how the change in [CO_2_] and the consequent FITC loading evoked by two different durations of stimulation (1 and 5 min) varies with the V_max_ of CAII. Reduction from the control value (15 mM/s) to 1 mM/s simulates the effect of acetazolamide, whereas an increase of V_max_ to 30 mM/s simulates application of L-Phe. D) A summary graph showing how FITC loading varies with stimulus duration with the 3 different values for the V_Max_ of CAII.

We added a rate of CO_2_ production that was proportional to mitochondrial ATP production, endowed the “paranode” with a K^+^ channel to give it a resting potential, Cx32 and carbonic anhydrase. The CO_2_ sensitive gating of Cx32 was based on the published CO_2_ dose response curves (Huckstepp et al., 2010). FITC was assumed to only permeate open Cx32 hemichannels and its transmembrane concentrations were calculated according to the GHK equation assuming that FITC had a net negative charge of -1. CA activity was modelled with Michaelis-Menten kinetics with K_M_ being based on literature values for CAII. The V_max_ of CA was a free variable that could be altered to mimic the effect of inhibition or allosteric enhancement of CA.

We altered the duration of electrical stimulation of the “paranode” from 1 to 20 mins and calculated the amount of dye loading. With a V_max_ of 15 mM/s, this gave a graph that was very similar to the experimentally obtained data (Fig 10D, compare to Fig 6B). To simulate the effect of acetazolamide we reduced the V_max_ of CA to 1 mM/s, and found an enhancement of dye loading that was once again very similar to the experimentally observed enhancement (Fig 10D, compare to Fig 6B). L-Phe can enhance the activity of CA by up to 3-fold. We found that increasing the V_max_ of CA twofold to 30 mM/s gave a very substantial reduction of dye loading that was similar to the experimentally observed effect of L-Phe (Fig 10D, compare to Fig 6B).

Our simplified model of the paranode suggests that CA is a key regulator of the local PCO_2_ and hence Cx32 gating. We also observed that when inhibition of CA was simulated by a reduction of V_max_ to 1 mM/s, the concentration of CO_2_ increased to a steady state value of 0.48 mM and there was a steady increase in FITC loading reaching a concentration of 0.7 µM after 30 minutes. Under the “control” conditions [CO_2_] had a steady state value of 0.05 mM and the FITC concentration after 30 minutes was only 20 pM. There is some support for this prediction of the model as we observed that acetazolamide did indeed increase the background FITC loading of nerve fibres by a small but significant amount (Fig 10, figure supplement 1).

The Matsuda model explicitly incorporates mitochondrial priming by electrical activity, and its use in our model reproduces the experimentally observed dye loading. This suggests that mitochondrial priming might also occur in the node/paranode, although this remains to be tested directly.

### Activity dependent entry of Ca^2+^ into the paranode is CO_2_ dependent

Our evidence so far supports the hypothesis that Cx32 is gated during action potential propagation by activity dependent generation of CO_2_ at the node. During electrical activity Ca^2+^ accumulates in the paranode (Lev-Ram and Ellisman, 1995). As we have previously shown Cx32 to be Ca^2+^ permeable (Butler and Dale, 2023), we tested whether this increase in paranodal Ca^2+^ could be caused by entry via the CO_2_-dependent opening of Cx32.

To measure intracellular Ca^2+^ we loaded isolated mouse sciatic nerve with Fluo4-AM. We found that exposure of the nerve to hypercapnic aCSF (70 mmHg) increased Fluo4 fluorescence in paranode-like structures indicating an increase in intracellular Ca^2+^ (Fig 11). The CO_2_ evoked increases in Fluo4 fluorescence were blocked by carbenoxolone indicating that they were channel mediated most likely via Cx32 (Fig 11, p = 0.0087 CBX compared to control).

**Figure 11:**
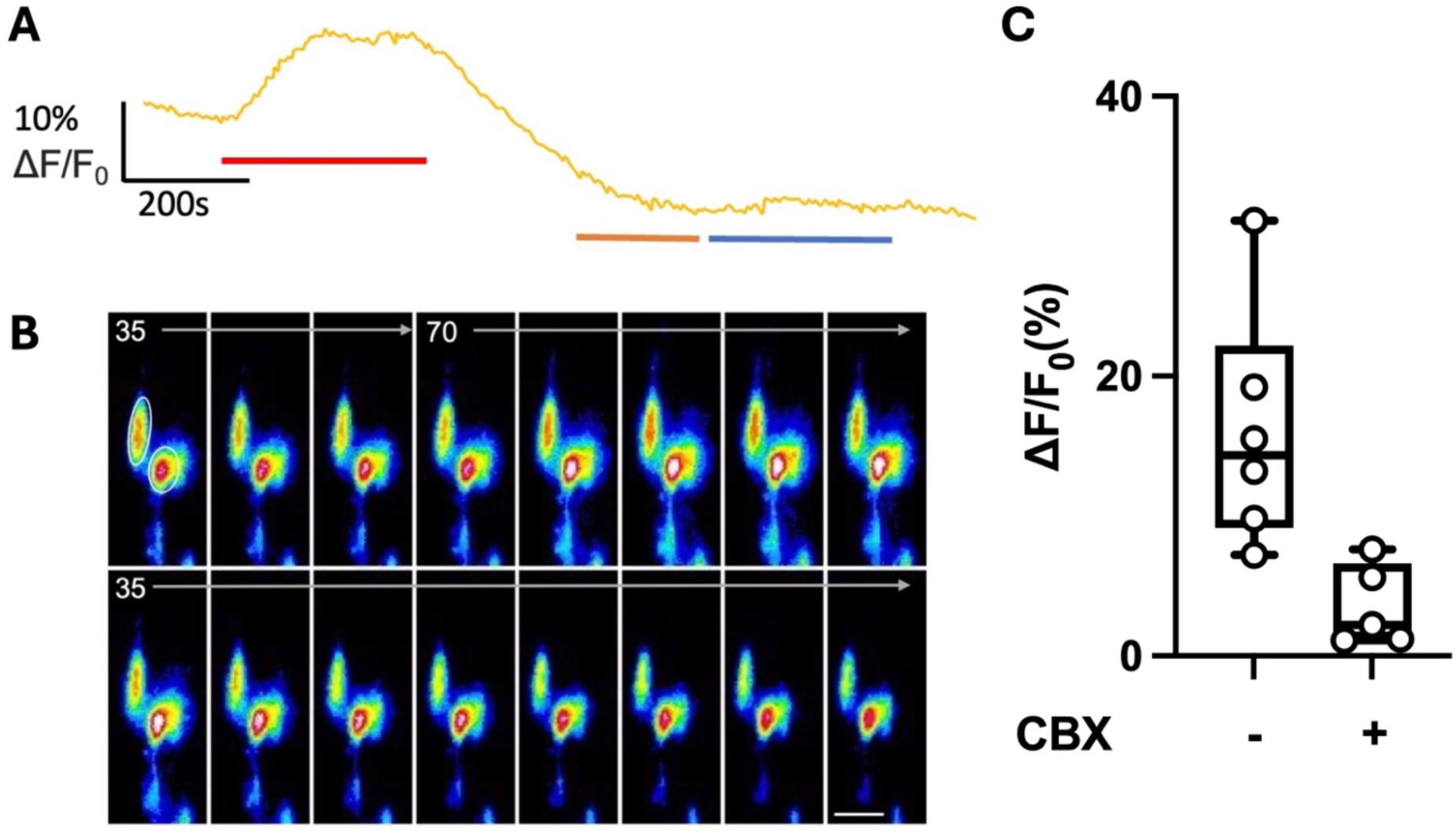
CO_2_ dependent Ca^2+^ influxes into Schwann cells are hemichannel dependent. A) Representative trace showing change in normalised Fluo4 fluorescence in response to 70mmHg aCSF (red bar), 35mmHg aCSF with the non-specific hemichannel blocker carbenoxolone (100 µM, orange bar) and 70 mmHg aCSF plus 100 µM carbenoxolone (blue bar).B) Representative images showing changes Fluo4 fluorescence in response to hypercapnic aCSF. The circles in the first panel show the measurement ROIs drawn around the paranodes. Scale bar = 10 µm. C) Boxplot showing the change in normalised fluorescence (ΔF/F_0_) in Fluo4 loaded Schwann cell paranodes evoked by 70 mmHg aCSF in the presence and absence of 100 µM carbenoxolone (CBX). Each datapoint consists of a paranode, with all the data collected from 4 sciatic nerves. Control vs CBX, MW test, p = 0.0087.

Having established the existence of CO_2_ dependent Ca^2+^ entry into the paranode, we next determined whether we could observe Ca^2+^ entry into the paranode during electrical stimulation and whether this was also CO_2_ dependent. To measure intracellular Ca^2+^ we expressed GCaMP8 under the control of the Mpz promoter to ensure Schwann cell specific expression (Fig 12A). Stimulation of the isolated sciatic nerve evoked increases in intracellular Ca^2+^ as reported by GCaMP8 fluorescence that could be enhanced by AZ and blocked by TC AQP1-1 (Fig 12 B-D). Use of Fluo4 loaded sciatic nerves replicated these data and showed an increase of Ca^2+^ into Schwann cell paranodes during electrical stimulation (Fig 12, figure supplement 1). Crucially, these transient increases depended upon CO_2_ production: they were significantly enhanced by acetazolamide (p<0.0001) and reduced by block of AQP1 by TC AQP1-1 (p<0.0001; Fig 12, figure supplement 1). Thus, the Ca^2+^ entry into the paranode during electrical stimulation depends on CO_2_ generated by the axon entering the paranode and most likely opening Cx32. This is consistent with earlier reports that show that Ca^2+^ accumulation in the paranode requires extracellular Ca^2+^ (Lev-Ram and Ellisman, 1995).

**Figure 12:**
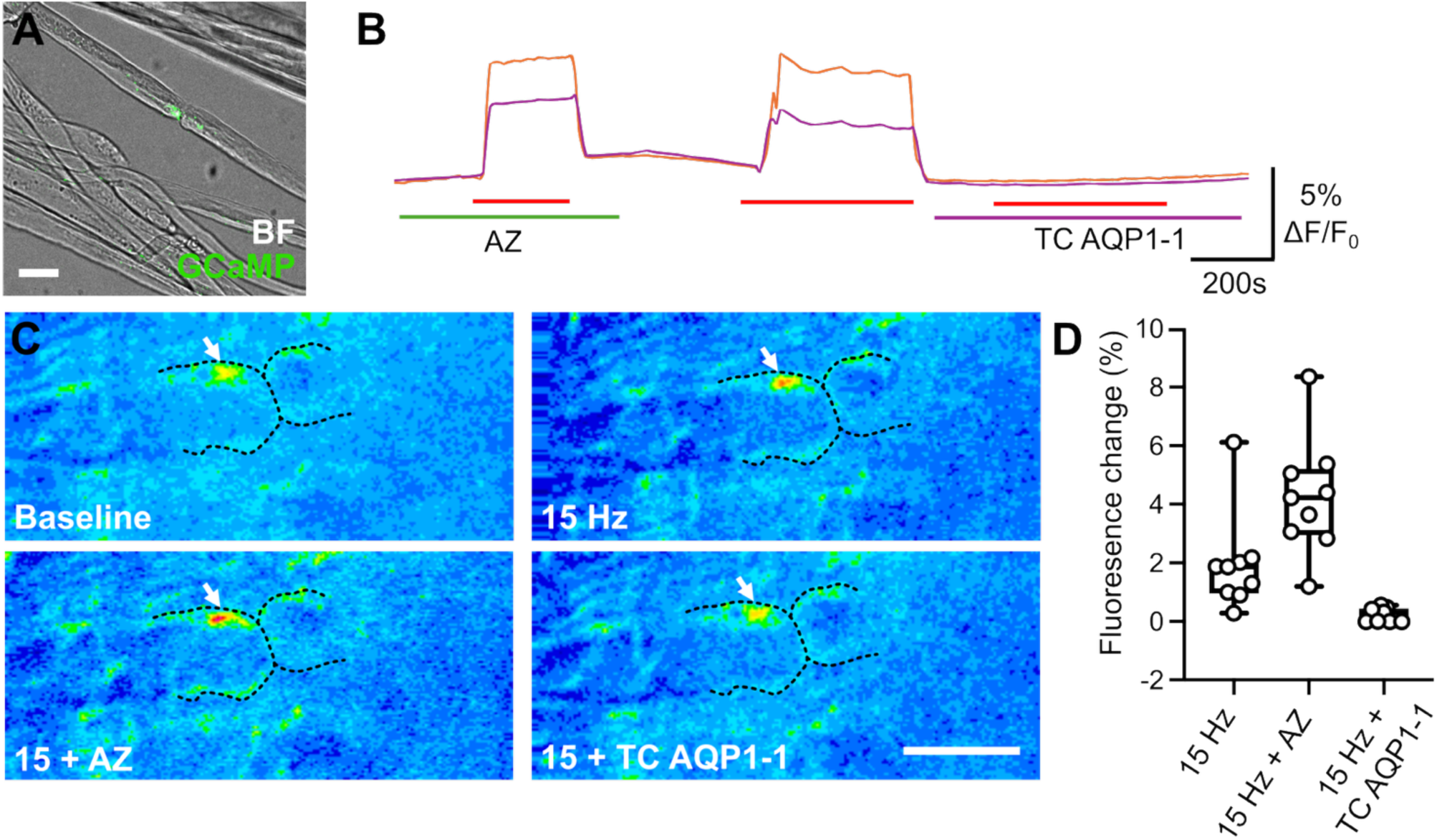
Activity dependent increase of intracellular Ca^2+^ in Schwann cell paranodes. A) Superimposed brightfield (BF) and fluorescence image of GCaMP8 transduced nerve. To show expression at the paranode. B) Representative GCaMP8 traces showing change in normalised fluorescence in response to 15 Hz electrical stimulation (red bar) in the presence of acetazolamide (AZ, green bar) or TC-AQP1-1 (purple bar). C) The fluorescence images show before (Baseline), during stimulation of the nerve (15 Hz), during stimulation in presence of AZ (15 + Az) and stimulation in the presence of TC AQP1-1 (15 + TC AQP1-1). Scale bar = 5 µm; black dashed line indicates the shape of an individual fibre, white arrow indicates hotspot of GCaMP8 fluorescence at the paranode. D) boxplot showing the change in normalised GCaMP8. fluorescence (ΔF/F_0_) evoked Schwann cell paranodes in response to 15 Hz electrical stimulation in the control, with AZ and with TC AQP1-1. Kruskal Wallis ANOVA, p < 0.0001. Pairwise MW comparisons: control vs AZ, p = 0.0142; control vs TC AQP1-1, p<0.0001. Each datapoint consists of a paranode, with all the data collected from 4 sciaSc nerves.

### Activity dependent slowing of conduction velocity is CO_2_ dependent

We observed that hypercapnic aCSF, FCCP and electrical activity consistently induced FITC loading into the outer myelin layer, suggesting the occurrence of CO_2_ dependent gating of Cx32 in this outermost membrane. Were this to occur, it should increase the leakage of current across the myelin sheath. Saltatory conduction depends on local current circuits travelling down the core of the axon to depolarise that next node (Huxley and Stampfli, 1949). If more current were to leak through the sheath before reaching the next node there should be a small but measurable slowing of conduction velocity (Huxley and Stampfli, 1949; Bakiri et al., 2011). We would therefore predict that during more intense electrical activity in nerve, there should be more CO_2_ production and thus a slowing of conduction velocity.

To test this, we measured the CAP firstly under low frequency stimulation (1 Hz), exposed the nerve to a period of high frequency stimulation (15 Hz for 10 mins, to elevate local PCO_2_) and then remeasured the CAP under low frequency stimulation (1 Hz). We found high frequency stimulation increased the delay from the stimulus artefact to the peak of the CAP by 0.11 ms (median, 95% CI: 0.04 to 0.17) (Fig 13). To demonstrate that this slowing was CO_2_-dependent we manipulated the components of the CO_2_ signalling system. 100 µM acetazolamide significantly increased the delay to the peak of the CAP caused the high frequency stimulation (p = 0.0016, Fig 13). Conversely, 1 mM L-Phe or 80 µM TC AQP1-1 reduced the effect of high frequency stimulation on the delay to the peak of the CAP (respectively p = 0.0317 and p = 0.0079, Fig 13). Note that once again, TC AQP1-1 had no effect on the amplitude of the CAP (Fig 13D).

**Figure 13.**
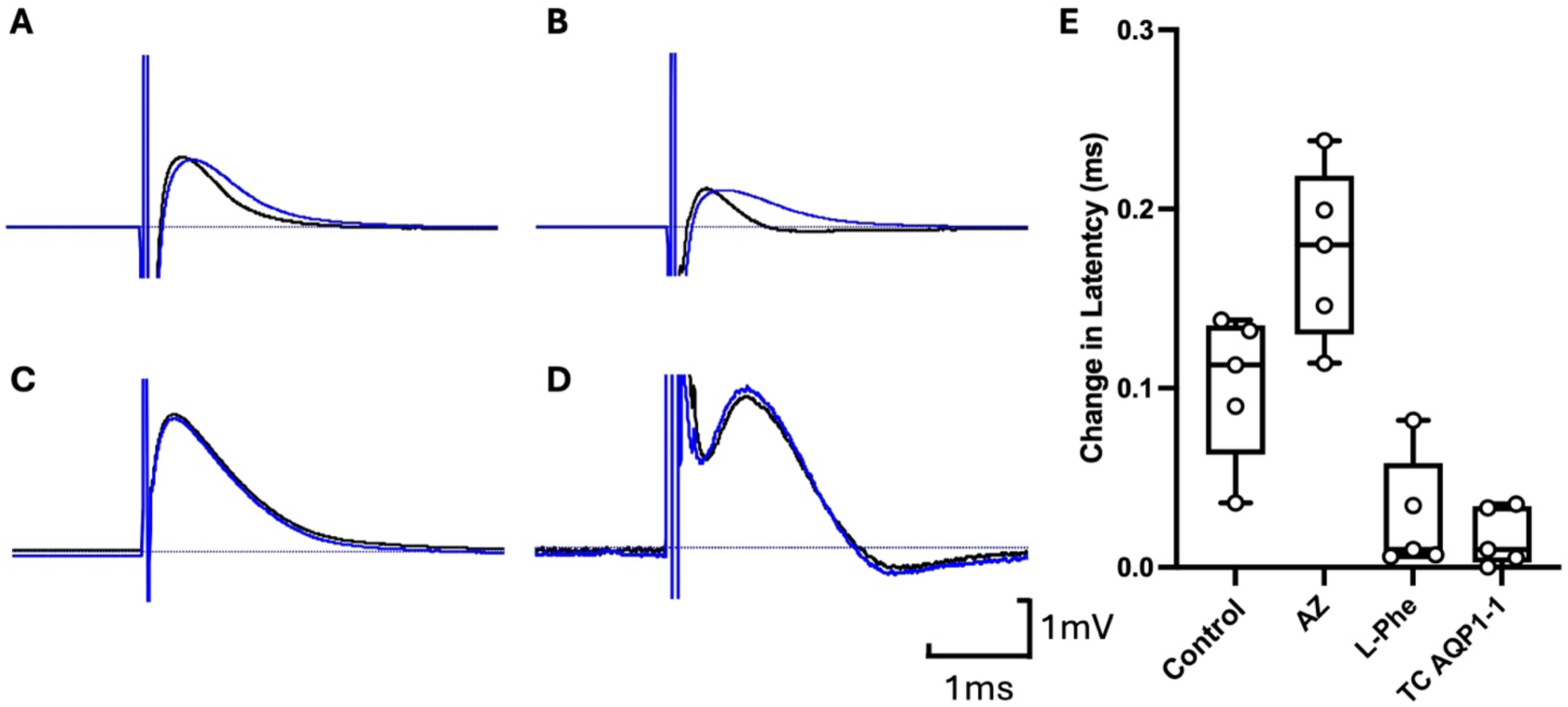
CO_2_ dependent slowing of conduction velocity following high frequency stimulation. The CAP was evoked at 1 Hz and then 10 mins of 30 Hz stimulation was given prior to remeasuring the CAP at 1 Hz stimulation frequency. Representative CAPs from mouse sciatic nerve prior to high frequency stimulation (Black trace), and after (Blue trace) for WT nerves: A) in the absence of any compound; B) with 100 µM acetazolamide; C) with 1 mM L-Phenylalanine;D) with 80 µM TC AQP1-1. E) Boxplot showing the change in latency (time to peak of CAP) before and after high frequency stimulation for Control, Acetazolamide (AZ), L-Phe and TC AQP1-1. Kruskal Wallis ANOVA p=0.0016. Pairwise MW tests: control vs AZ, p=0.0317; control vs L-Phe, p=0.0159; control vs TC AQP1-1, p=0.0079.

We noticed that the period of high frequency stimulation broadened the CAP and slightly reduced its amplitude. This was particularly exaggerated in the presence of acetazolamide (Fig 13, fig supplement 1). To understand this effect on the shape of the CAP, we made a simple model of the CAP based upon 2000 individual axons each having an identically shaped action potential. To reflect the distribution of fibre diameters reported in sciatic nerve (Assaf et al., 2008) the conduction velocities were given a normal distribution skewed to lower velocities (Fig 13, fig supplement 1). The CAP was simply the sum of all of the individual action potentials. We then slowed the velocity of each fibre by the same proportion and computed the CAP for different amounts of slowing and calculated the 10-90% rise time, the time to peak, and peak amplitude of the CAP (Fig 13, fig supplement 1). This showed that under these simplified assumptions, changes in the shape of the CAP of the type we observed experimentally would be expected from slowing the conduction velocities in all fibres by the same proportion.

## Discussion

### Activity dependent gating of Cx32

In this paper we have investigated the mechanism of activity dependent Cx32 hemichannel gating in peripheral myelin. Previously, the opening of Cx32 has been posited to depend on either its intrinsic voltage sensitivity (Abrams et al., 2002) or as a downstream consequence of an increase in cytosolic Ca^2+^ within the paranode (Stauch et al., 2012; Carrer et al., 2018). Here, we have tested an alternative hypothesis: that cell-to-cell signalling mediated via CO_2_ produced in the axon is the primary trigger for Cx32 gating in the paranode. Our hypothesis explicitly links Cx32 opening in the paranode to the energetic demands of action potential propagation in the node.

In our experiments, we assessed Cx32 gating via entry of the membrane impermeant dye, FITC. Our results support our new hypothesis in several respects. Firstly, Cx32 gating in response to axon stimulation was greatly reduced by blocking AQP1, which is CO_2_ permeable (Endeward et al., 2006; Musa-Aziz et al., 2009; Michenkova et al., 2021) and thus provides a route for passage of CO_2_ from axon to paranode. The inhibition of AQP1 had no effect on the CAP, eliminating a possible alternative interpretation that blocking this channel directly affected NaV1.8 (Zhang and Verkman, 2010).

Secondly, inhibition of CA with acetazolamide greatly increased the activity dependent gating of Cx32. Thirdly, facilitation of CA activity by applying an allosteric enhancer, L-Phe, greatly reduced activity dependent gating of Cx32. Fourthly, the effect of FCCP showed that mitochondrially generated CO_2_ was sufficient to gate Cx32. Fifthly, our use of low doses of H_2_O_2_ to block aconitase greatly reduced activity-dependent dye loading, indicating its dependence on a functioning Krebs cycle.

Finally, application of GDPβS to block all GPCR based signalling had no effect on activity-dependent gating of Cx32. Together these results suggest that CO_2_ is acting as a cell-to-cell signal and is the prime trigger for Cx32 opening during action potential propagation. In the light of these results, it is interesting that elasmobranchs, the first vertebrates to evolve a fully myelinated nervous system (Salzer and Zalc, 2016), have an orthologue of Cx32 that has identical CO_2_ sensitivity to human Cx32 (Dospinescu et al., 2019).

As there are no selective pharmacological blockers for Cx32, our evidence that Cx32 is the conduit for activity and CO_2_ dependent FITC loading into the paranode is indirect. Nevertheless, our combined evidence is compelling for the following reasons. Cx32 is the only known large-pored channel expressed in Schwann cells that is directly sensitive to gaseous CO_2_. We know that FITC permeates Cx32 and the CO_2_ dose dependence of FITC loading matches that of Cx32. We have eliminated both Cx31.3 (not CO_2_ sensitive) and TRPA1 (unaffected by a selective blocker of this channel) as the conduit. However, the strongest evidence to support our hypothesis is that expression of Cx32^DN^ in the Schwann cell blocks activity dependent FITC loading. This shows that dye loading depends upon both Cx32 and CO_2_ binding to Cx32.

### Possible localisation of the components required for CO_2_ signalling and their relation to the energetics of action potential generation

Our data and previously published studies (Toews et al., 2007) support the localisation of Cx32 in the paranode and outer myelin layers. AQP1 also localises to both the paranode and axon either in the node or close to the node in the paranodal region of the axon. Nevertheless, Cx32 and AQP1 are not restricted to these locations and are found for example in the internodal regions. Colocalisation analysis (restricted to the paranode/nodal regions) shows that in these regions Cx32 and AQP1 show significant proximity as do AQP1 and a mitochondrial marker CytC. Cx32 also shows significant colocalisation with CytC. Mitochondria are thus likely to be present in both the paranode and node. This is consistent with other studies suggesting mitochondrial localisation to the axonal node (Zhang et al., 2010; van Hameren et al., 2019). It should be noted that mitochondria are not restricted to the axonal node (Zhang et al., 2010).

In myelinated axons, the voltage-gated Na^+^ influx and K^+^ efflux occurs at the node of Ranvier. The transmembrane ionic gradients at the node need to be restored via the actions of Na^+^-K^+^ ATPases. The nodes of Ranvier are likely therefore to be the major sites of ATP generation and consumption and thus the production of CO_2_. However, as we cannot directly measure CO_2_ production, the site of its production remains a matter of supposition.

The binding site for CO_2_ on Cx32 is intracellular and CO_2_ must therefore cross both the nodal and paranodal membranes. The localisation of the key components (Cx32, AQP1, mitochondria and CA) in the nodal/paranodal region will shorten the diffusion path between the source of CO_2_ (mitochondria) and its ultimate target Cx32. This would potentially speed the dynamics of the CO_2_ signal. CA, which provides an efficient removal mechanism for CO_2_, shows restricted localisation to the paranode. This supports our hypothesis of CO_2_ entry through paranodal AQP1.

### The important roles of AQP1 and carbonic anhydrase

Whilst there has been controversy over the role of CO_2_ permeable channels in enabling transmembrane CO_2_ fluxes, it is now accepted that biological membranes are only poorly permeable to CO_2_ and a channel mediated mechanism is required (Boron et al., 2011). Our data further support this idea, as blockade of APQ1 prevents the activity dependent gating of Cx32. Given that our data also show that CA activity limits the gating of Cx32, the colocalization of AQP1 and Cx32 may be important. As AQP1 will be the entry point for CO_2_ into the paranode, its colocalization with Cx32 may favour CO_2_ binding to Cx32 over capture and conversion to carbonic acid by CA.

Our simplified model of the paranode sheds further light on the regulation of CO_2_ signalling and the gating of Cx32. The Matsuda model (Matsuda et al., 2020) incorporates priming of mitochondrial ATP production by electrical activity and hence this is also implicit in our paranode model. Mitochondrial priming will make CO_2_ production more rapid than if it depended on ATP depletion to occur first. This implies that CO_2_ production could vary relatively quickly with activity patterns and thus report the dynamics of action potential firing. It will be important to directly test this prediction by measuring the dynamics of mitochondrial ATP production in the node relative to imposed electrical activity.

Our model also predicts that complete inhibition of CA will give some Cx32 gating under non-stimulated conditions. We observed increased baseline loading of FITC into the nerves during acetazolamide giving support for this prediction. This suggests that an important role of CA is to prevent basal rates of CO_2_ production being sufficient to gate Cx32. Our model also suggests that CA activity regulates the extent to which activity dependent production of CO_2_ can gate Cx32. The effects of L-Phe and acetazolamide lend support to this prediction. Thus, the model suggests that CA controls the dynamic range of the CO_2_ signal and is likely to be an important regulator of CO_2_ mediated signalling. By keeping paranodal cytosolic PCO_2_ low, CA not only reduces the basal gating of Cx32 but importantly also maintains a concentration gradient that favours entry of CO_2_ into the paranode.

### Physiological consequences of CO_2_ dependent signalling

We have further shown that two other aspects of Schwann cell physiology and function depend on the CO_2_-dependent gating of Cx32. Firstly, the well documented increase of intracellular Ca^2+^ into the paranode evoked by nerve stimulation appears to largely depend on the opening of Cx32 and can be modified in the same way as dye loading by manipulating CA activity or blocking AQP1. Secondly, given the localisation of Cx32 in the outer myelin layer and the observation of activity- and CO_2_-dependent dye loading into that outer layer, we hypothesised that there should be activity dependent slowing of nerve conduction. This is because any opening of Cx32 hemichannels should reduce the resistance to current flow across the myelin sheath. We did indeed observe a small degree of activity dependent slowing of nerve conduction velocity. Crucially, this too depended upon CO_2_ production and could be altered by manipulating CA activity or blocking AQP1.

### Links to CMTX

Given our evidence suggests that the CO_2_ sensitivity of Cx32 is critical for its gating during action potential propagation, we might expect that any mutations that affect this sensitivity could precipitate CMTX. We have previously examined the effect of 14 CMTX mutations on the CO_2_ sensitivity of Cx32 (Butler and Dale, 2023). We found that 5 completely removed its CO_2_ sensitivity, 3 greatly reduced its sensitivity while the remainder had no apparent effect. It should be noted that two mutations of K124 (Bone et al., 1997; Fattahi et al., 2017) and K104 (Williams et al., 1999; Wang et al., 2015), the critical residues for detection of CO_2_ (K104E, K104T, K124E and K124N, not included in our published study), have also been identified as possible CMTX mutations.

These results, while supportive of our hypothesis, are not conclusive as several of the 8 CMTX mutations that altered CO_2_ sensitivity have been documented to also affect other facets of channel function. However, the CMTX mutation E102G stands out as causing moderate severity CMTX while still permitting the formation of gap junction channels and hemichannels with apparently normal voltage dependence and ATP permeability. Because E102G involves the loss of CO_2_ sensitivity of the hemichannel in the absence of other known functional effects on the hemichannel (Abrams et al., 2003) it lends some support to the hypothesis that the CO_2_-dependence of Cx32 may be important for the health of myelin and that its loss could precipitate CMTX. Further exploration of CO_2_- and Cx32-dependent signalling in myelin may suggest new strategies to treat peripheral neuropathies and peripheral nerve injury,

## Methods

All experiments were performed in accordance the United Kingdom Home Office Animals (Scientific Procedures) Act (1986) with project approval from the University of Warwick’s AWERB and licence PP7458325.

### Sciatic nerve isolation

All mice used were C57BL/6, aged at least 6 weeks. Sciatic nerves were isolated following the protocol described in Rich et al., 2018. Placing a few drops of ice cold aCSF loosened the perineural membrane allowing its easy removal with forceps, beginning at one cut end of the nerve, and moving inward. Once the perineural membrane was removed, forceps were carefully placed in between the seams of the larger bundle of fibres before being teased apart with careful lateral movement. Removal of the perineural membrane and slight teasing was sufficient to obtain dye loading, with extensive dissection to small bundles or individual fibres only occurring post-fixation for immunohistochemistry and visualisation.

### Immunocytochemistry

Antibodies used:

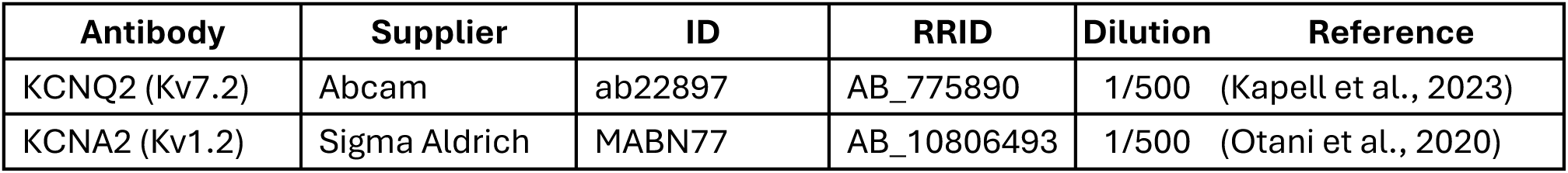
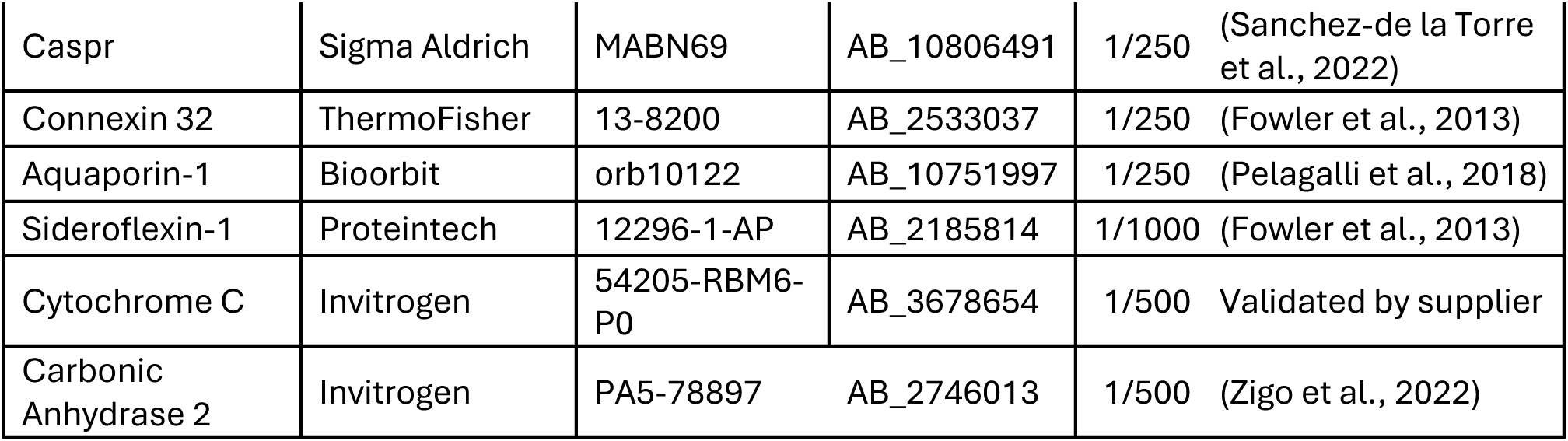

Sciatic nerves were first washed with PBS three times, before being fixed in 4% PFA for 45 mins. Nerves were then washed in PBS three times and blocked using PBS containing 4% BSA and 0.1% Triton X-100 for 24 hr. Nerves were teased prior to immunostaining. Primary antibody was diluted in PBS containing 4% BSA and 0.1% Triton X-100 and added to nerves and left to incubate, constantly shaking, for 48hrs at 4℃. Nerves were then washed using PBS containing 0.1% Triton X-100 six times at 10 min intervals. The appropriate secondary antibodies diluted in PBS containing 4% BSA and 0.1% Triton X-100 and added to coverslips and left to incubate, constantly moving, for 2.5 h. The secondary antibody was washed using PBS containing 0.1% Triton X-100 six times at 10 min intervals. Nerves were again blocked for 24hrs. To co-stain with a further primary antibody from the same species, antibody conjugation was used (ProteinTech FlexAble corallite®). Conjugated antibodies were diluted in PBS containing 4% BSA and 0.1% Triton X-100 and added to nerves and left to incubate, constantly moving, for 48 hr at 4℃. The conjugated antibody was washed using PBS containing 0.1% Triton X-100 six times at 10 min intervals. Nerves were then placed onto a glass slide and further dissected using the tips of hypodermic syringes, yielding individual nerve fibres. Nerves where then mounted using Fluorshield^™^ with DAPI mounting medium (Sigma-Aldrich, Cat# F6057), placing a glass coverslip on top. Nerve fibres were subsequently imaged using the Zeiss-880 and Zeiss 980 confocal LSMs, specifically using the 488, 561 and 630 nm lasers. FIJI software was used for further analysis. Images were also taken on a Nikon N-SIM S with dual camera, utilising a 100X oil immersion lens. 470, 561 and 640 nm lasers were used.

### Solutions used

#### Control (35 mmHg PCO_2_) aCSF

124 mM NaCl, 3 mM KCl, 2 mM CaCl_2_, 26 mM NaHCO_3_, 1.25 mM NaH_2_PO_4_, 1 mM MgSO_4_, 10 mM D-glucose saturated with 95% O_2_/5% CO_2_, pH 7.4.

#### Hypercapnic (70 mmHg) aCSF

73 mM NaCl, 3 mM KCl, 2 mM CaCl_2_, 80 mM NaHCO_3_, 1.25 mM NaH_2_PO_4_, 1 mM MgSO_4_, 10 mM D-glucose, saturated with ∼12% CO_2_ (with the balance being O_2_) to give a pH of 7.4.

#### Depolarising (35 mmHg PCO_2_) aCSF

77 mM NaCl, 50 mM KCl, 2 mM CaCl_2_, 26 mM NaHCO_3_, 1.25 mM NaH_2_PO_4_, 1 mM MgSO_4_, 10 mM D-glucose saturated with 95% O_2_/5% CO_2_, pH 7.4.

### Pharmacological agents

| Compound | Supplier | Concentration (µM) |
| --- | --- | --- |
| Acetazolamide | Sigma Aldrich A6011 | 100 |
| Ammonium chloride | Invitrogen A15000.0B | 100 |
| GDPβS | Sigma Aldrich G7637 | 100 |
| L-Phenylalanine | Sigma Aldrich P2126 | 1000 |
| TC AQP1-1 | Tocris 5412 | 80 |

### Dye loading assay

Isolated nerves were obtained as described above and slightly teased apart with sharp needles as this was found to produce more profound and reliable dye-loading.

#### CO_2_-dependent dye loading

Isolated sciatic nerves were first washed in control aCSF before being superfused with either 35 mmHg aCSF or 70 mmHg aCSF containing 50 μM fluorescein isothiocyanate (FITC) for 10 minutes. To induce endogenous CO_2_ production via mitochondrial uncoupling isolated nerves were superfused with 35mmHg aCSF containing 10 μM FCCP (APExBIO) for 10 minutes. Following this, FITC was washed off by superfusion with 35 mmHg aCSF with no dye for 10 minutes. The nerves were then transferred through a series of vessels with 35mmHg aCSF to remove any remaining FITC.

#### Dye loading triggered by electrical stimulation

Nerves were pre-incubated in 35 mmHg aCSF, and any desired pharmacological agent, for 10 minutes prior to recording. Polished glass suction electrodes wrapped with a silver wire and backfilled with aCSF were used for stimulation and recording. The ends of nerves were gently sucked up into the suction electrodes such that orthodromic recordings were made, described in (Rich et al., 2018).

The recordings of the stimulus evoked CAPs were controlled by a National Instruments A/D interface (Model PCIe 6321) using the Strathclyde electrophysiology software program, WinWCP. A stimulator, Digitimer model DS3, was used to stimulate the nerve. The signal was amplified x1000 by an A-M Systems Inc Model 3000 AC/DC differential amplifier (A-M Systems, Sequim, WA 98382, USA). The signal was filtered at 20 kHz and 1 Hz and acquired at 20 kHz. To assess the validity of the CAP, the nerve was crushed between forceps at the conclusion of the experiment, leaving only the transient artefact.

Once a recording of the CAP had been successfully established, nerves were exposed to FITC during electrical stimulation at (30 Hz) of different durations (1-10 minutes). They were then washed to remove FITC as described above. As each mouse possesses 2 sciatic nerves, when pharmacological agents were used, one nerve from each animal would be stimulated in the absence of any pharmacological agents as a matched control. I-V curves were recorded prior to drug pre-incubation, during pre-incubation and following stimulation, to assess any effects of the used compound on the CAP. From the CAP traces rise time, rate of rise and latency could be calculated using WinWCP.

#### Fixation and imaging of dye-loaded nerves

Nerves were then fixed using 4% PFA for 45 minutes. Nerves were subsequently imaged using Zeiss 880 or 980 confocal LSMs, specifically using the 488, 561 and 633nm lasers. FIJI software was used for further analysis. The statistical replicate was a single region of interest (ROI) and these were obtained from 5 nerves for each condition.

### Measurement of intracellular pH with BCECF

Mouse sciatic nerve was dissected as previously described. BCECF-AM dissolved in DMSO was diluted into 35 mmHg aCSF to a final concentration of 2.5 μM. A hypodermic needle was blunted and joined to a fine glass capillary via a short length of tubing. Etched tungsten wire was used to make a small incision in the middle on the nerve, from which the nerve was teased open slightly. Whilst holding the incision open the capillary loaded with BCECF was inserted and injected. The nerve was placed into 35mmHg aCSF to wash for 3 minutes. The nerve was then placed into a recording chamber, immobilised with a platinum wire harp and superfused with 35 mmHg aCSF. The BCECF-loaded nerves were imaged by epifluorescence (Scientifica Slice Scope, Cairn Research OptoLED illumination, 60x water Olympus immersion objective, NA 1.0, Hamamatsu ImagEM EM-SSC camera, Metafluor software). BCECF was excited using 470nm LED, with fluorescent emission being recorded every 4 seconds between 507 and 543nm. Once a stable fluorescence baseline was reached, the various test solutions were superfused onto the nerve. Intracellular pH was then calibrated using Nigericin (James-Kracke, 1992).

The statistical replicate was a single nerve and 5 nerves were recorded for each condition.

### Adeno-associated viral (AAV) constructs used

All AAVs were commercially produced (brainVTA) using the AAV9 serotype. To direct gene expression to Schwann cells, a minimal P0 (Mpz-mini) promoter was used (Sargiannidou et al., 2015; Kagiava et al., 2021; Georgiou et al., 2023). GCaMP8 was tagged with an LCK sequence, tethering it to the inner Schwann cell membrane. Cx32^DN^ (Butler and Dale, 2023) was flanked by IRES-mCherry (van de Wiel et al., 2020), giving cytoplasmic mCherry in transduced Schwann cells. Aliquots were stored at -80 until used.

### Transduction of sciatic nerve *in vivo*

Surgical procedures were performed under the authority of the UK Home Office Licence PP7458325. Anaesthesia was induced by inhalation of isoflurane (4%; Piramal Healthcare Ltd, Mumbai, India) in pure oxygen (4 L·min^-1^). The mouse was then placed on a temperature regulated heating pad (TCAT-2LV, Physitemp) to maintain body temperature. A face mask was used to maintain anaesthesia (isoflurane, intranasal, 0.5–2.5% in pure oxygen1 L·min^-1^) throughout the surgery. Atropine was provided (subcutaneous, 0.05 mg/kg) before surgery to stop pleural effusion. Adequacy of anaesthesia was assessed by respiratory rate, body temperature, and pedal withdrawal reflex. Preoperative meloxicam (subcutaneous, 2 mg/kg) and postoperative buprenorphine (subcutaneous, 0.05 mg/kg) were provided for analgesia. Unilateral intraneural injection into the sciatic nerve were performed on six- to eight-week-old male C57BL/6 mice, using 600-900 nL of AAV. The injections were performed manually, using a graduated micropipette attached to 1ml syringe, at a rate ∼200 nl/min. The micropipette was left in the nerve for 5 minutes following injection before removal. If any animal showed signs of pain in the days following surgery, additional analgesia (oral meloxicam) was administered as required. Postoperatively, 4-5 weeks were allowed to achieve maximal AAV expression, before dissection, electrophysiology and imaging as previously described.

### Measurement of intracellular Ca^2+^ with GCaMP8 or FLuo4

Nerves were placed into the recording chamber and anchored with a platinum harp. The nerve was perfused with 35 mmHg aCSF until a stable baseline was reached. The desired solution was then perfused until a stable level had been reached before being washed.

For experiments that utilised Fluo4 imaging, Fluo4-AM dissolved in Pluronic™ F-127 (Thermo Fisher Scientific P3000MP) with constant sonication and vortexing and was diluted into 35mmHg aCSF to a final concentration of 2.5 μM. Nerves were then incubated for 20 minutes before being washed in 35 mmHg aCSF.

To enable simultaneous electrical stimulation and imaging of GCaMP8 or Fluo4 fluorescence, the nerves were mounted between electrodes within a bespoke micro-perfusion chamber constructed of Sylgard™-184. Proprietary software was used to control nerve stimulation at 15Hz, record the CAP and perform offline analysis.

Nerves were imaged by epifluorescence (Scientifica Slice Scope, Cairn Research OptoLED illumination, 60x water Olympus immersion objective, NA 1.0, Hamamatsu ImagEM EM-SSC camera, Metafluor software). GCaMP8 or Fluo4 were excited with a 470nm LED, and fluorescent emission between 507 and 543nm recorded every 4 seconds.

The statistical replicate was a single ROI (paranode) and these were obtained from 5 nerves for each condition.

### Measurement of activity dependent conduction velocity slowing

Using isolated nerves, CAPs were recorded as described above using the Strathclyde electrophysiology software. The baseline of CAP was recorded at low frequency (1Hz) for 30 seconds. High frequency stimulation (30 Hz) was applied for 10 minutes. Following this the nerves were then stimulated for 30 seconds at 1Hz. The CAPs from before and after high frequency stimulation were averaged and compared. A recording from an isolated nerve was considered as a statistical replicate.

### Cell culture and transfection

The Cx31.3 gene sequence were synthesized by IDT and subcloned into the pCAG-GS-mCherry vector. DNA gBlock was amplified using PCR with primers (IDT) Plasmids were generated using Gibson assembly. The presence of the correct assembly was confirmed by DNA sequencing (GATC biotech). The Cx31.3 construct was inserted upstream of mCherry, with a short 12 amino acid linker (GVPRARDPPVAT).

pDisplay-GRAB_ATP1.0-IRES-mCherry-CAAX was a gift from Yulong Li (Addgene plasmid # 167582 ; http://n2t.net/addgene:167582 ; RRID:Addgene_167582)

Parental HeLa DH cells were grown in Low-glucose Dulbecco’s modified eagle medium (DMEM) supplemented with 10% FBS and 50μg/mL penicillin/streptomycin. The HeLa DH cells were plated onto coverslips at a density of 7.5 X 10^4^ cells per well of a 6 well plate and transiently transfected using a mixture of 1 μg each of the Cx31.3 construct and GRAB_ATP_ and 3 μg PEI for 6hrs. Cells were imaged 48 hours after transfection. We used a protocol to measure ATP release from cells developed and described in our previous work (Butler and Dale, 2023).

### Analysis of GRAB_ATP_ fluorescence

Analysis of GRAB_ATP_ was performed in ImageJ (Schneider et al., 2012). Cell recordings were corrected for any motion using the Image Stabilizer plugin (Li, 2008). For cells expressing both Cx31.3 and GRAB_ATP_, an ROI was drawn around the GRAB_ATP_ expression and median fluorescence measured for each image. The fluorescence pixel intensity (F) was normalised to the baseline fluorescence (F_0_). The change in normalised fluorescence (ΔF/F_0_) evoked each stimulus, CO_2_ and 50 mM KCl, was recorded for each cell.

We converted changes in normalised fluorescence evoked by 70 mmHg pCO_2_ and 50 mM KCl into the concentration of ATP released by normalising them to the ΔF/F_0_ produced by a 3 μM ATP calibration solution. Over this range, the calibration curve for GRAB_ATP_ is approximately linear (Wu et al., 2022; Butler and Dale, 2023). Statistical comparisons were performed considering each cell as an independent replicate. Five transfections were performed.

### Analysis of Immunohistochemical colocalization

The JACOP plugin (Bolte and Cordelieres, 2006) was used to calculate the Manders’ coefficients M1 and M2. The convention we have used throughout the paper is that for colocalization of A with B, M1 represents the proportion of A pixels that overlap with B pixels, and M2 would represent the proportion of B pixels overlapping with A pixels. Colocalisation analysis was restricted to the nodal/paranodal regions.

As a control, one channel was rotated 90^0^ and analysis was re-run using the same thresholds. Each datapoint represents an ROI, or a paranode, with all data points coming from at least 5 nerves.

### Modelling of paranode

The paranode was modelled as a cell with a mitochondrion, a K^+^ leak channel, Cx32 and CA. Mitochondrial ATP production was modelled as per Matsuda et al (Matsuda et al., 2020). This involves a time-dependent variable Y, that stimulates ATP production, where X is a variable that is proportional to the duration of electrical stimulation, Y_0_ is the steady state value of Y and relaxes to that value with a time constant of τ_1_. K_d1_ and n_1_ are parameters for the Hill equation that determines how variable X alters the value of Y:

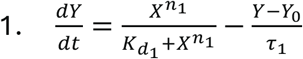

A time-dependent variable Z determines the use of ATP:

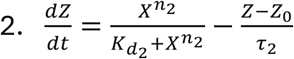

where Z_0_ is the steady state value of Z and relaxes to that value with a time constant of τ_2_ and K_d2_ and n_2_ are parameters for the Hill equation that determine how variable X alters the value of Z.

The rate of change of ATP concentration is thus:

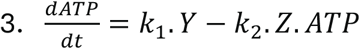

Where k_1_ and k_2_ are rate constants for synthesis and breakdown of ATP respectively.

To adapt this model to the paranode we first defined the rate of CO_2_ production is proportional to ATP production, and used the Michaelis Menten equation to calculate CO_2_ conversion to carbonic acid:

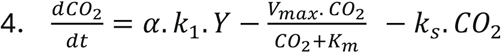

Where V_max_ and K_m_ are the maximal velocity of CA and affinity of CA for CO_2_ respectively, α is a rate constant for the production of CO_2_. For completeness, we also allowed for spontaneous conversion of CO_2_ to carbonic acid -determined by the first order rate constant, k_s_. Variables Y and k_1_ have the same meaning as in equation 3.

The rate of membrane potential change of the paranode was calculated from:

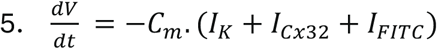

Where C_m_ is the whole cell membrane capacitance, I_K_ the K^+^ leak current, I_Cx32_ the current through Cx32 and I_FITC_ the current carried by FITC.

I_K_ is described by the Goldman Hodgkin Katz (GHK) equation:

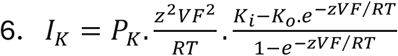

Where P_K_ is the maximal whole cell permeability to K^+^, z the valence. R is the universal gas constant, T the absolute temperature (in Kelvin) and F the Faraday constant. K_I_ and KO are respectively the intracellular and extracellular concentrations of K^+^.

I_Cx32_ is described by:

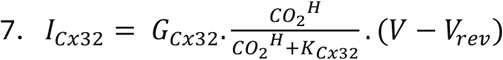

Where G_Cx32_ is the maximal whole cell conductance for Cx32, K_Cx32_ is the affinity of Cx32 for CO_2_, H is the Hill coefficient of CO_2_ binding and V_rev_ is the reversal potential of the current through Cx32.

I_FITC_ (through Cx32) is described by:

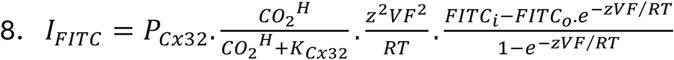

Where P_Cx32_ is the maximal whole cell permeability of Cx32 to FITC and z the valence of FITC. R is the universal gas constant, T the absolute temperature (in Kelvin) and F the Faraday constant. FITC_i_ and FITC_o_ are respectively the intracellular and extracellular concentrations of FITC. K_Cx32_ and H have the same meaning as in equation 7.

The rate of change of FITC_i_ with time is thus:

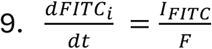

Where F is the Faraday constant.

These differential equations were coded with Matlab, and a 4^th^ order Runge-Kutta ODE solver with adaptive step size used to numerically integrate them and thus calculate the production of CO_2_ during electrical stimulation and the extent FITC loading. The Matlab code along with a command line interface is presented in the source files: CO2Cx32Plot.m and CO2Cx32Model.m.

### Modelling of CAP

Using Matlab, a compound action potential was computed from summing 2000 individual action potentials based on the product of two Boltzmann equations to give a realistic shape (compoundAP.m). The action potentials were given a random delay (representing conduction velocity) based on a mean ± SD described by a Gaussian distribution with a skew factor. This was chosen to reflect the skewed distribution of axon diameters in the sciatic nerve (Assaf et al., 2008). Slowing could be introduced by slowing the conduction velocity of every individual action potential by a fixed proportion of its delay.

### Statistical analysis

All quantitative data are presented as box and whisker plots where the box represents the interquartile range, the bar represents the median, and the whiskers represent 1.5 times the interquartile range, or the range if this is less. Individual data points are superimposed onto boxplots. Statistical analysis was via the Kruskal Wallis one-way ANOVA (KW test) followed by pairwise Mann Whitney U-tests with correction for multiple comparisons via the false discovery method (Curran-Everett, 2000) with the maximum rate of false discovery set at 0.05. For analysis of the GRAB_ATP_ recordings in which the CO_2_ and 50 mM KCl stimuli were applied to the same cell, these data were considered to be paired and comparisons of the amount of ATP released by each stimulus was therefore performed with the Wilcoxon Matched Pairs Signed Rank test. All pairwise tests were two sided and all calculations performed with GraphPad PRISM.

## Supporting information

Source data for Fig 2 and supplements

Source data for Fig 4 and supplements

Source data for Fig 6 and supplements

Source data for Fig 7 and supplements

Source data for Fig 8 and supplements

Source data for Fig 9 and supplements

Source data for Fig 10 and supplements

Source data for Fig 11 and supplements

Source data for Fig 12 and supplements

Source data for Fig 13 and supplements

Matlab code for paranode model

Matlab code to run paranode model

## Acknowledgements

We thank Dr Joao Correia, Institute of Microbiology & Infection, University of Birmingham, for assistance with the super resolution microscopy.

## Author Contributions

JB: Conceptualisation, data collection and curation, investigation, writing – original draft, formal analysis. LM: Investigation, data collection and curation. AB^1^: sciatic nerve transduction, training and review; AB^2^: Conceptualisation, training and review. ND: Conceptualisation, modelling, supervision, writing – original draft, review and editing. All authors reviewed the final draft.

## Data availability

All data is included in the supplementary files for each figure.

## Figures and legends

**Figure 2 figure supplement 1.**
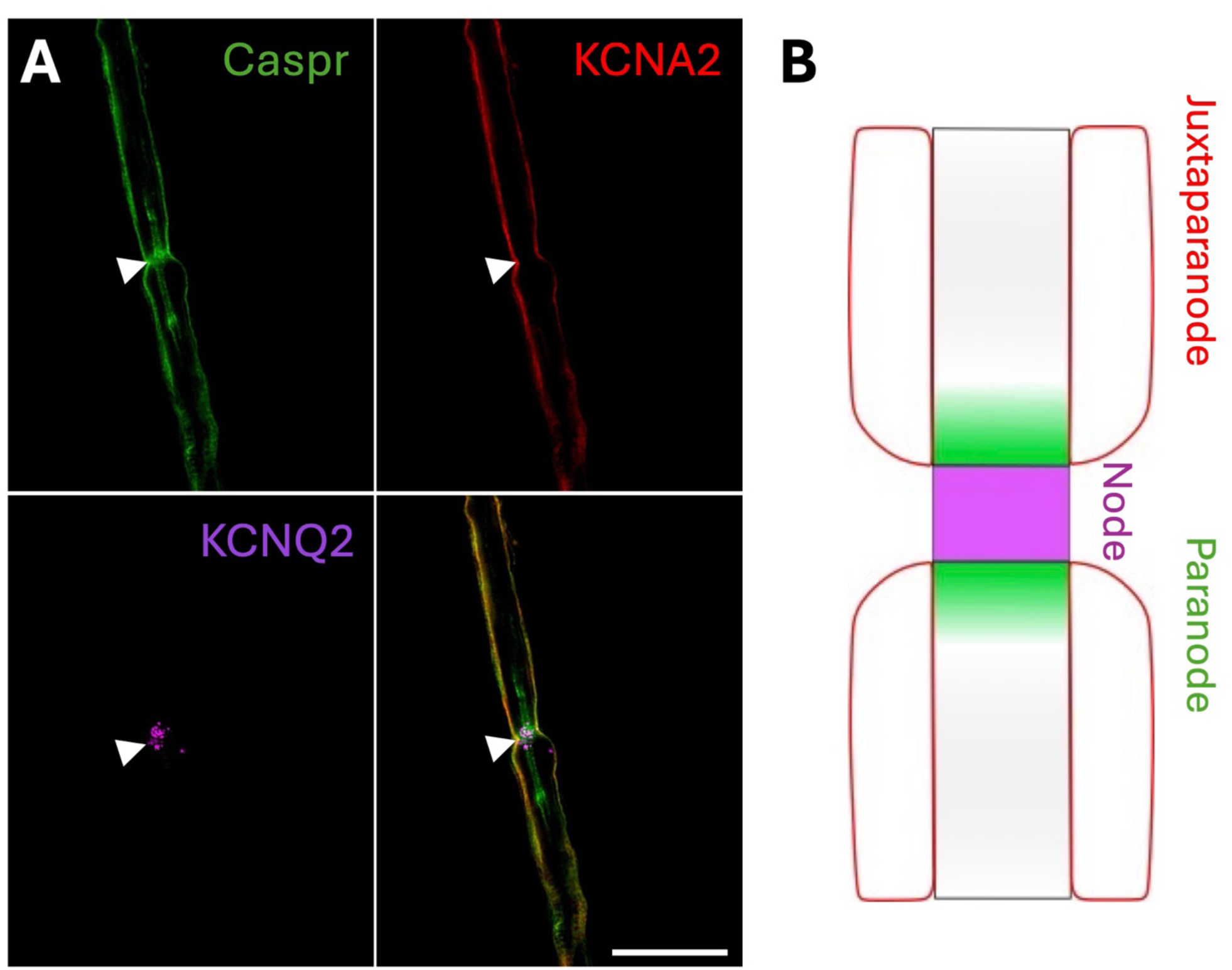
Markers for the node, paranode and juxtaparanode in the isolated mouse sciatic nerve. A) Location of the axonal node (KCNQ2) and Schwann cell paranode (Caspr) within an isolated mouse sciatic nerve fibre. KCNA2 is expressed in the outer myelin layer. Arrowheads indicate the node. Scale bar = 15 µm. Confocal LSM image, single optical plane. B) Schematic indicating the same locations within an isolated nerve fibre.

**Figure 2 figure supplement 2.**
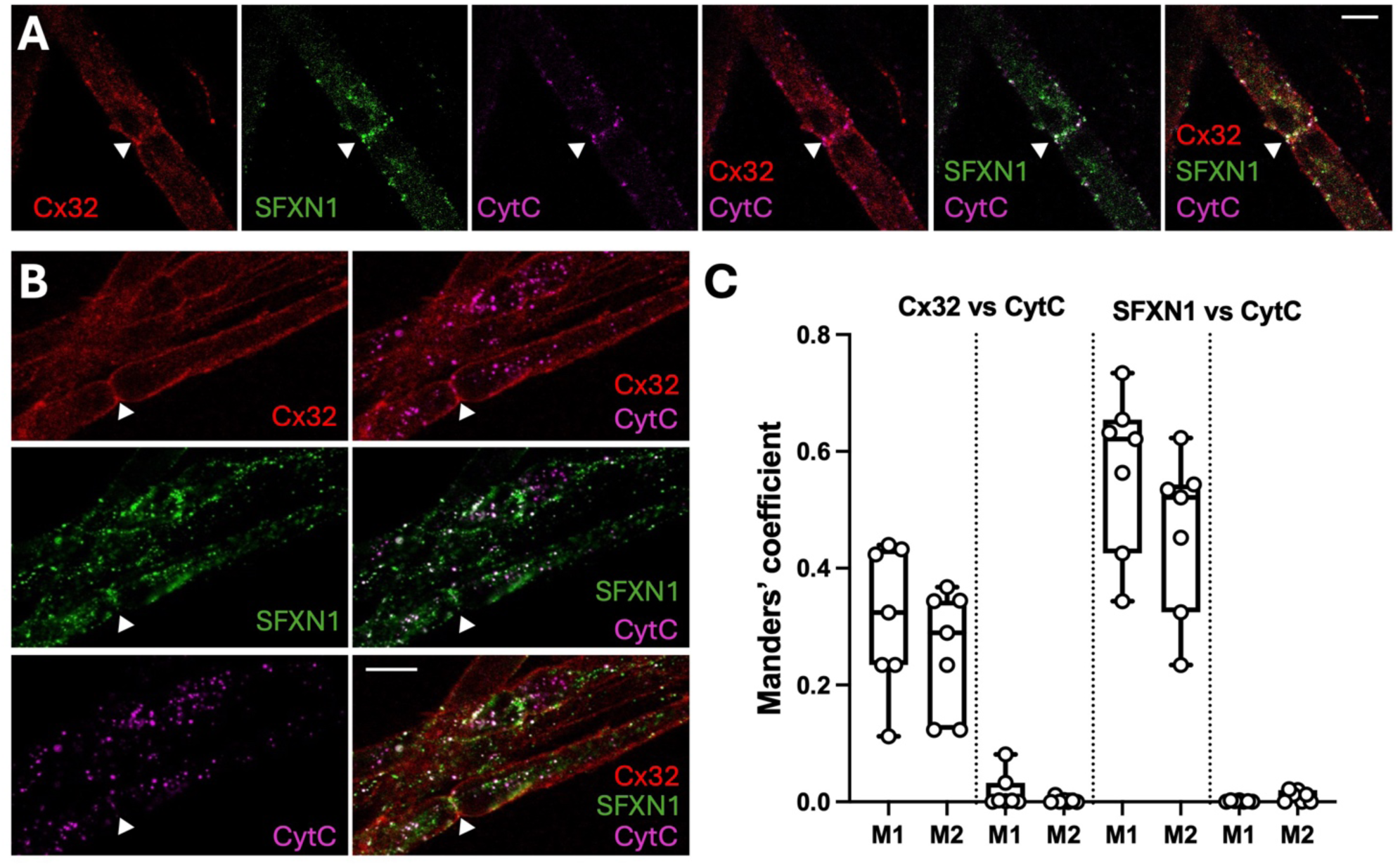
Cx32 colocalises with mitochondria in the Schwann cell paranode, alongside SFXN1. A, B) Representative SIM images in single optical plane showing the localisation of Cx32, CytC and SFXN1 in isolated mouse sciatic nerve. Arrowheads depict the node. Scale bars – 10μm. C) Boxplots showing degree of colocalization between Cx32 and CytC and SFXN1 and CytC. M1 is the proportion of Cx32 (or SFXN1) that colocalises with CytC and M2 is the reverse proportion of CytC that colocalises with Cx32 (or SFXN1). Control measurements (to right of dotted line for each pair) used these same images with one channel flipped 90° and the same thresholds as when measuring colocalization. Kruskal Wallis ANOVA: Cx32 vs CytC, p = 0.0001; SFXN1 vs CytC, p < 0.0001.

**Figure 2 figure supplement 3.**
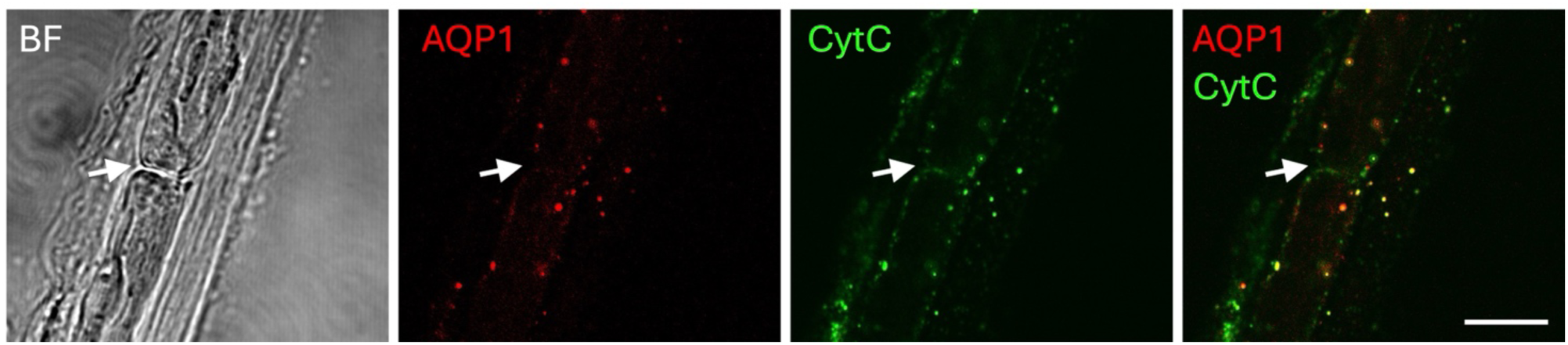
The localization of AQP1 in relation to mitochondria, CytC, in isolated mouse sciatic nerve. Representative SIM images in single optical plane showing the localisation of AQP1 and CytC in isolated mouse sciatic nerve. Arrows indicate the node. Scale bar – 10μm.

**Figure 4 figure supplement 1:**
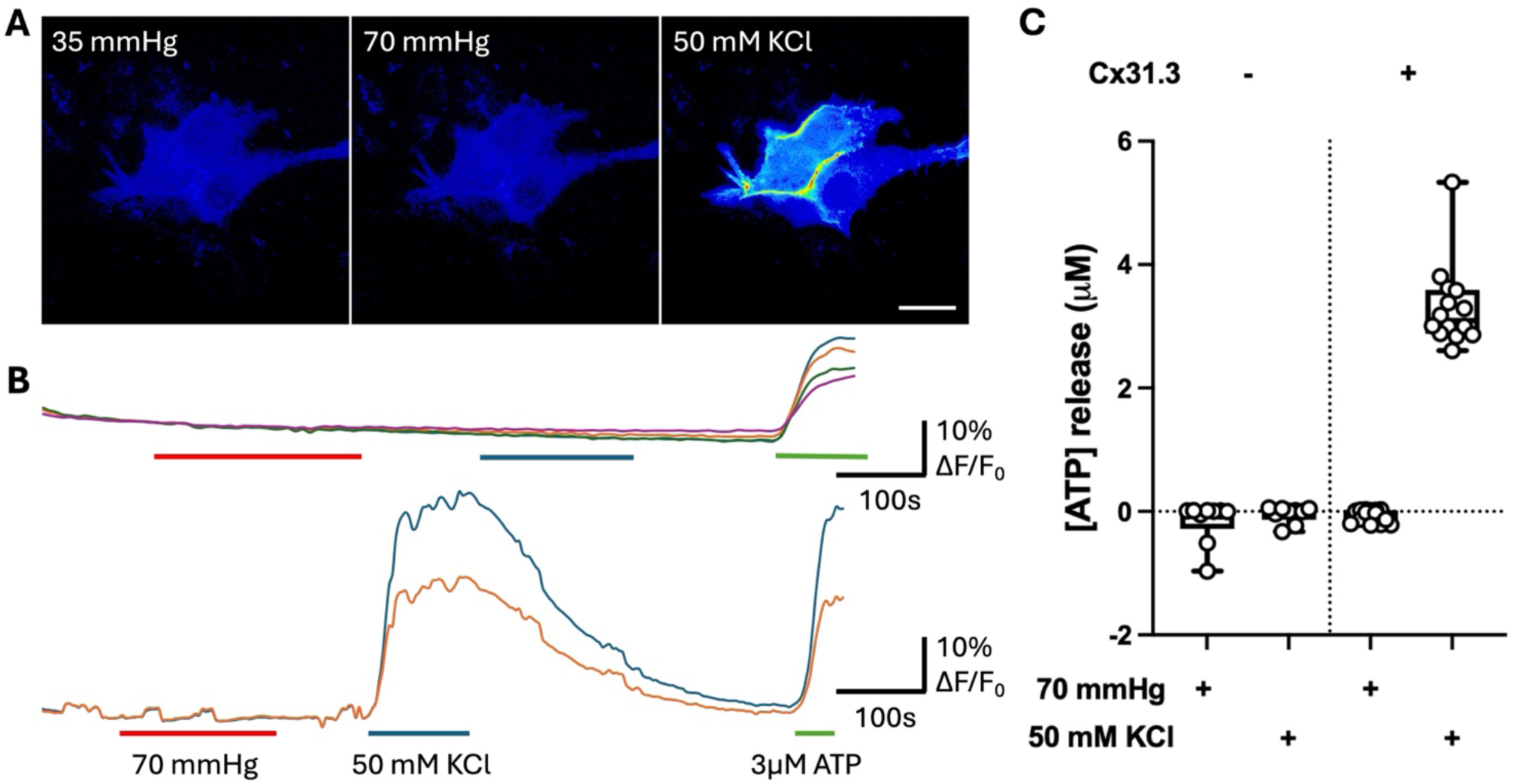
Cx31.3 does not open in response to hypercapnia. A) Representative images of GRAB_ATP_ fluorescence, encoded by a 16 colour LUT in Cx31.3 transfected HeLa cells in response to control (35 mmHg), hypercapnic (70 mmHg) and depolarising (50 mM KCl) aCSF. Scale bar 20 μm. B) Top, traces showing the normalised GRAB_ATP_ fluorescence for cells transfected only with GRAB_ATP_. Bottom, traces from the cells transfected with both Cx31.3 and GRAB_ATP_ as shown in (A). C) Summary statistic box plots showing the [ATP] release, calculated as a ratio from the normalised fluorescence change evoked by a certain solution or stimuli and the fluorescence change evoked by 3 µM ATP. Cells expressing only GRAB_ATP_ do not exhibit changes in fluorescence to 70 mmHg or 50 mM KCl aCSF. Each data point represents a cell, with all cells coming from at least three transfections.

**Figure 6 figure supplement 1.**
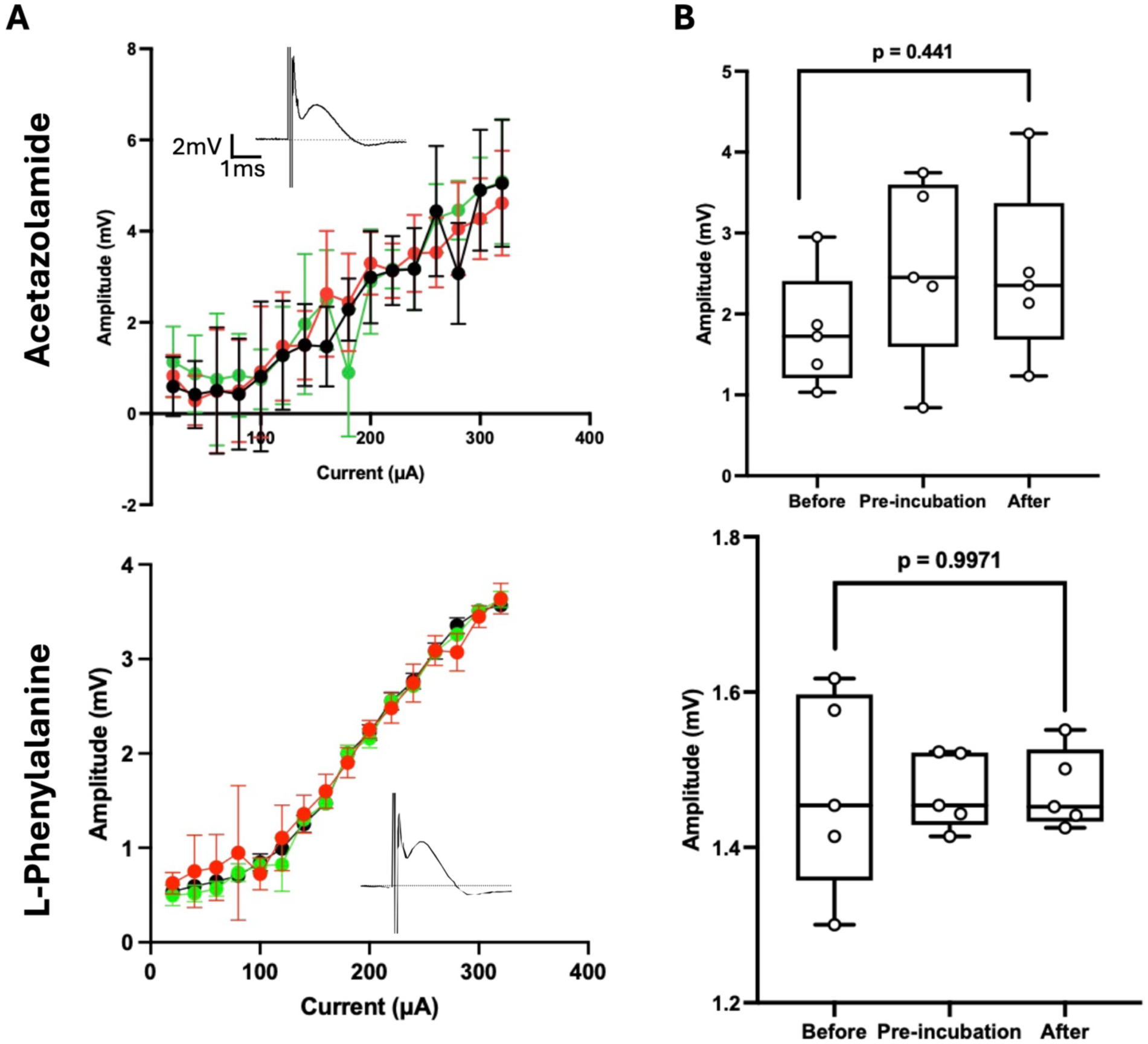
Acetazolamide and L-Phe do not alter the CAP. A) Current - voltage input-output curves for the CAP, comprising data from all nerves subjected to either 100 µM acetazolamide or 1 mM L-Phenylalanine. Curves were produced before application of the respective compound (green circles), at the end of the pre-incubation of the compound (red circles) and at the end of dye loading (black circles) to show that the compounds had no effect on the amplitude of the CAP. N=5 nerves for each condition. Insets show the averaged CAP during incubation with the compound. B) Boxplots showing the half maximum CAP amplitude for each respective condition and compounds laid out in (A). Kruskal Wallis ANOVA: Acetazolamide, p = 0.4441; L-Phenylalanine, p = 0.9917.

**Figure 7 figure supplement 1.**
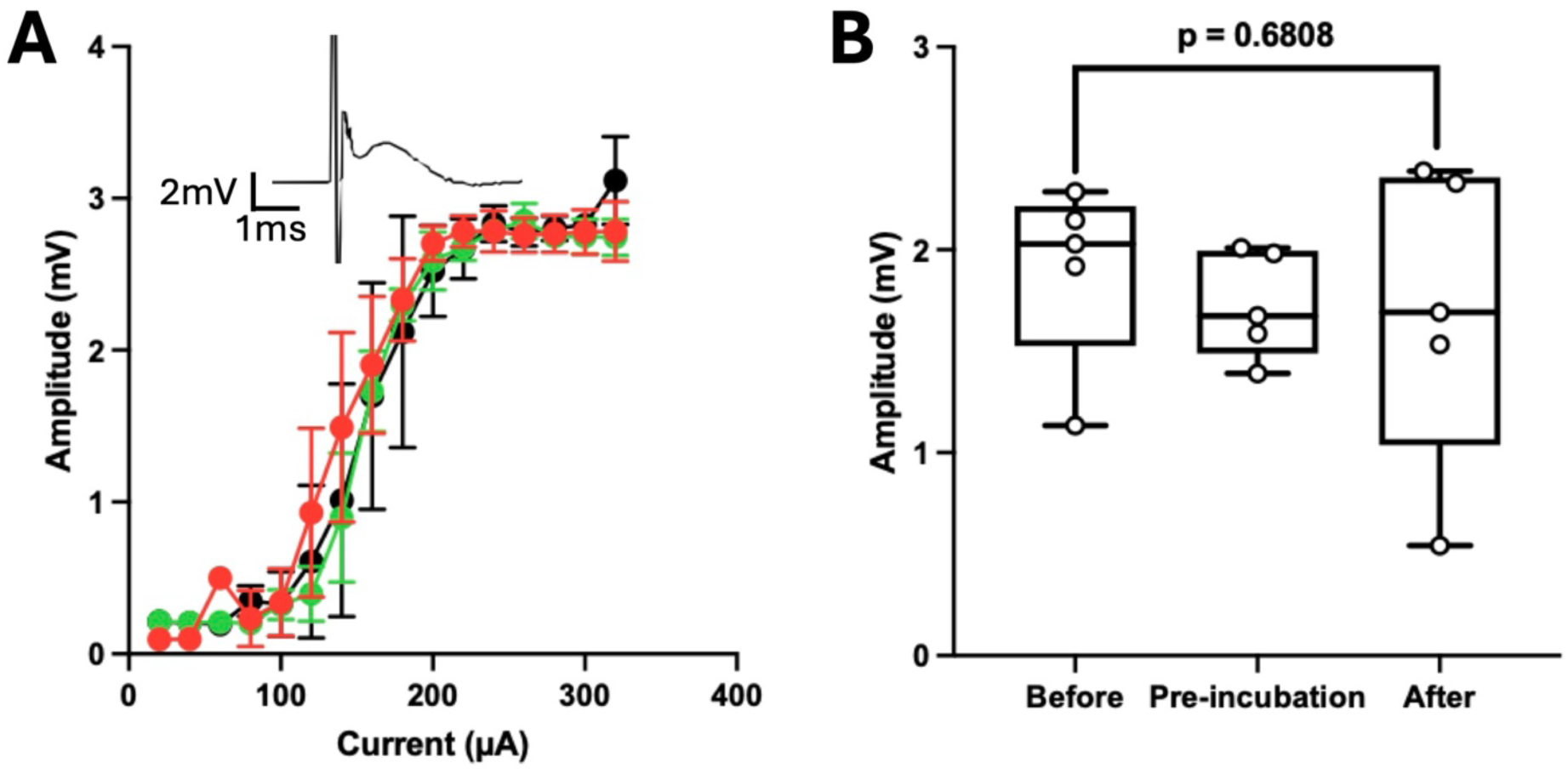
50 µM H_2_O_2_ does not alter the CAP. A) Current -voltage input-output curves for the CAP comprising data from all nerves subjected to 50 µM H_2_O_2_. Curves were produced before application of H_2_O_2_ (green circles), at the end of the pre-incubation (red circles) and at the end of dye loading (black circles) to show that H_2_O_2_ had no effect on the amplitude of the CAP. N=5 nerves for each condition. Inset shows the averaged CAP during incubation with the compound. B) Boxplots showing the half maximum CAP amplitude for each respective condition in (A). Kruskal Wallis ANOVA: p = 0.6808.

**Figure 8 figure supplement 1.**
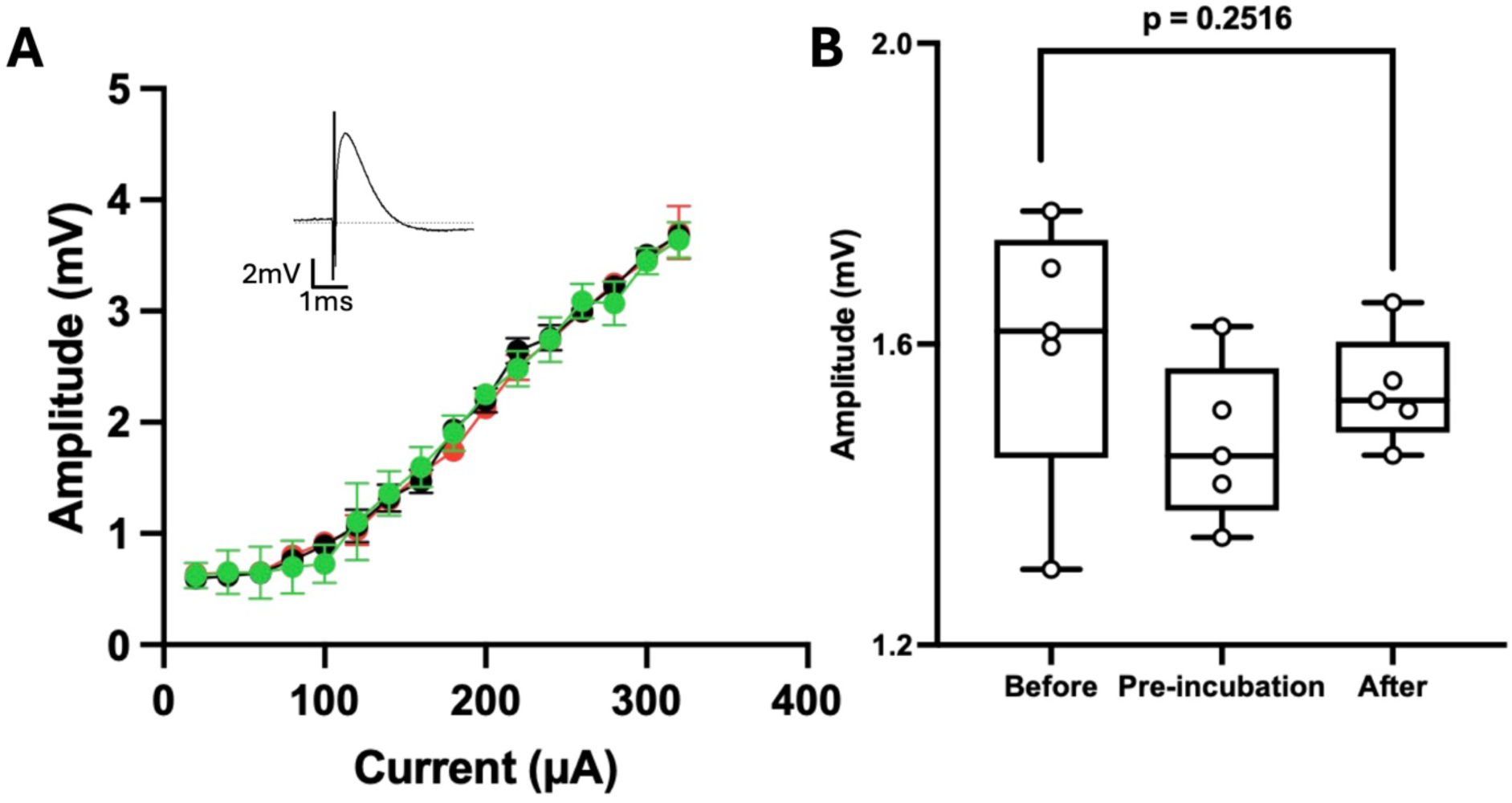
TC AQP1-1 does not alter the CAP. A) Current -voltage input-output curves for the CAP comprising data from all nerves subjected to 80 µM TC AQP1-1. Curves were produced before application of the TC AQP1-1 (green circles), at the end of the pre-incubation with TC AQP1-1 (red circles) and at the end of dye loading (black circles) to show that TC AQP 1-1 had no effect on the amplitude of the CAP. N=5 nerves for each condition. Inset shows averaged CAP during incubation with TC AQP1-1. B) Boxplots showing the half maximum CAP amplitude for each respective condition in (A). Kruskal Wallis ANOVA: p = 0.2516.

**Figure 8 figure supplement 2.**
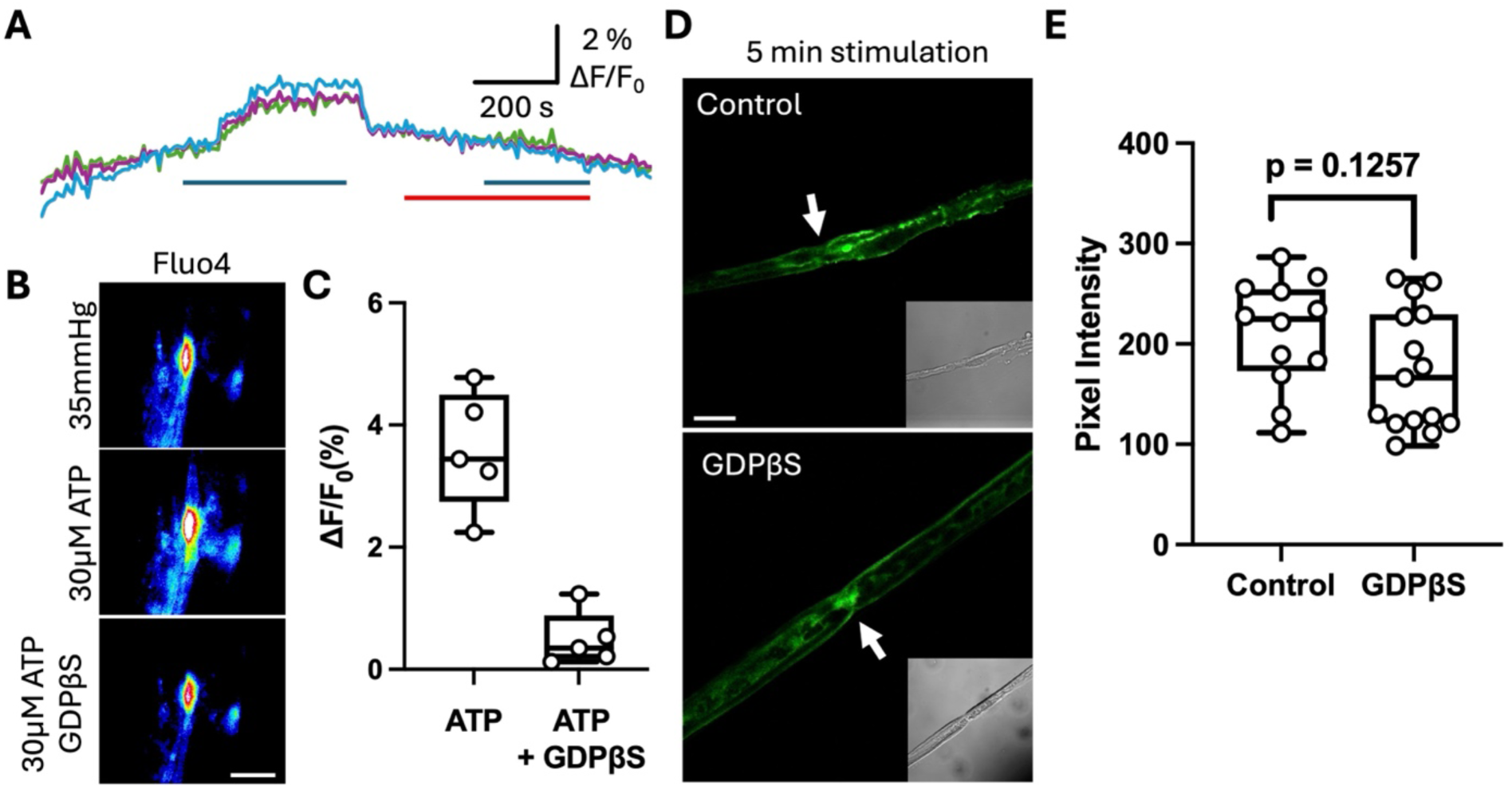
GDPβS blocks ATP-induced increases in paranodal Ca^2+^ but does not alter activity dependent FITC loading into the paranode. A) Recording of ATP-induced increase in intracellular Ca^2+^ at the paranode (blue bar). When ATP was applied after 100 µM GDPβS (red bar), no increase in Fluo4 fluorescence was seen. B) Images of Fluo4 fluorescence showing effect of ATP with and without GDPβS (same recording as A). Scale bar = 5 µm. C) Boxplots summarizing that GDPβS blocks the change in fluorescence induced by ATP. Each point is a single paranode from a single nerve. D) Images of FITC loading into sciatic nerve induced by 5 min electrical stimulation with and without 100 µM GDPβS. Arrows point to paranode, insets show corresponding brightfield images. Scale bar 10 µm. E) Boxplots showing that GDPβS has no effect on activity dependent (30 Hz stimulation) FITC loading. Each point represents a separate ROI from 5 different nerves.

**Figure 8 figure supplement 3.**
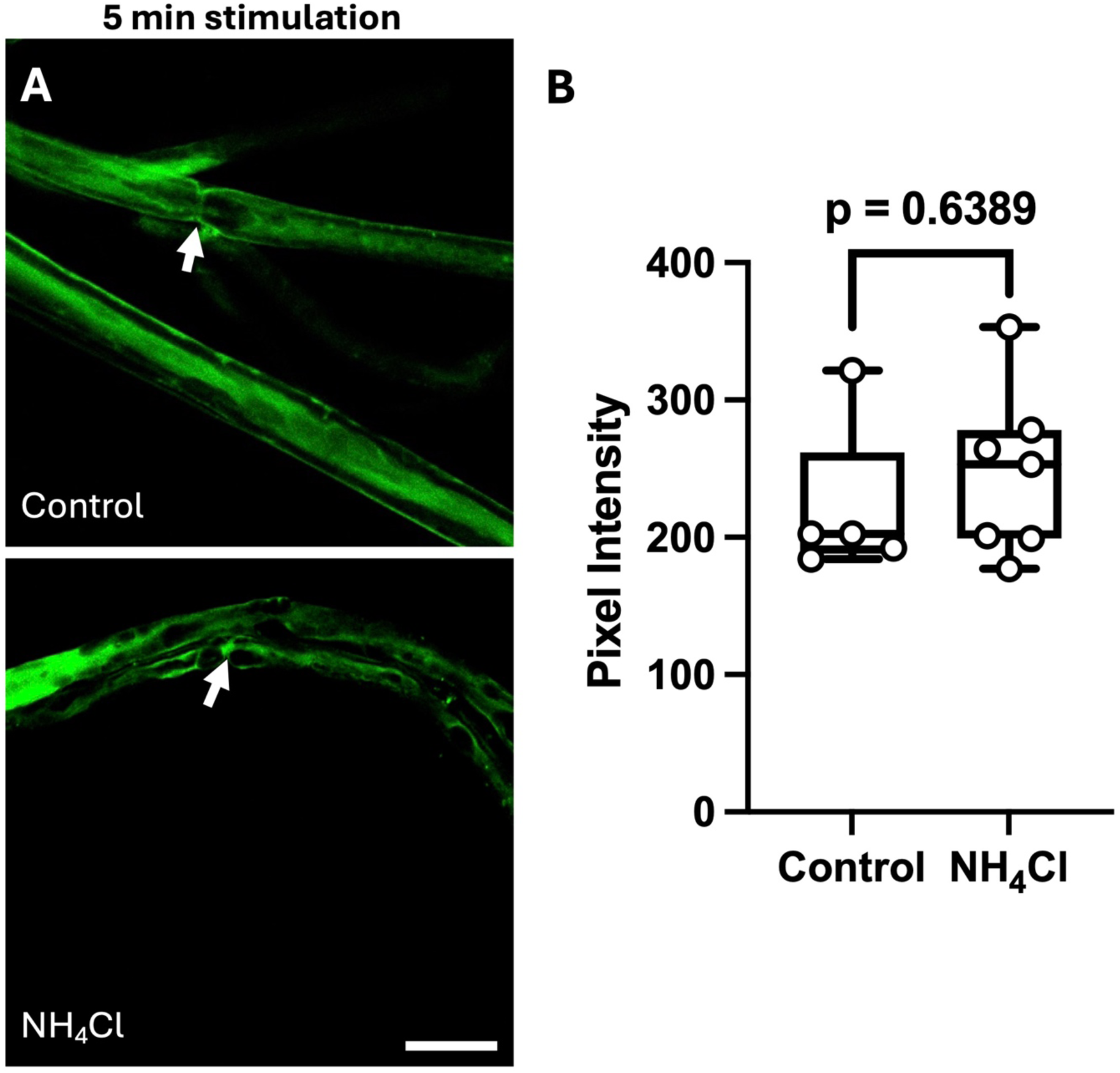
NH_4_Cl does not alter FITC loading into the paranode. A) Images of FITC fluorescence in isolated nerves stimulated for 5 minutes at 30 Hz, in the control (35 mmHg) and after treatment with 100 µM NH_4_Cl. Arrows indicate paranode. Scale bar 15 µm. B) Summary graph of FITC fluorescence in nerves in response to 5 minutes stimulation in each condition. Each point represents a separate ROI from 5 different nerves.

**Figure 8 figure supplement 4.**
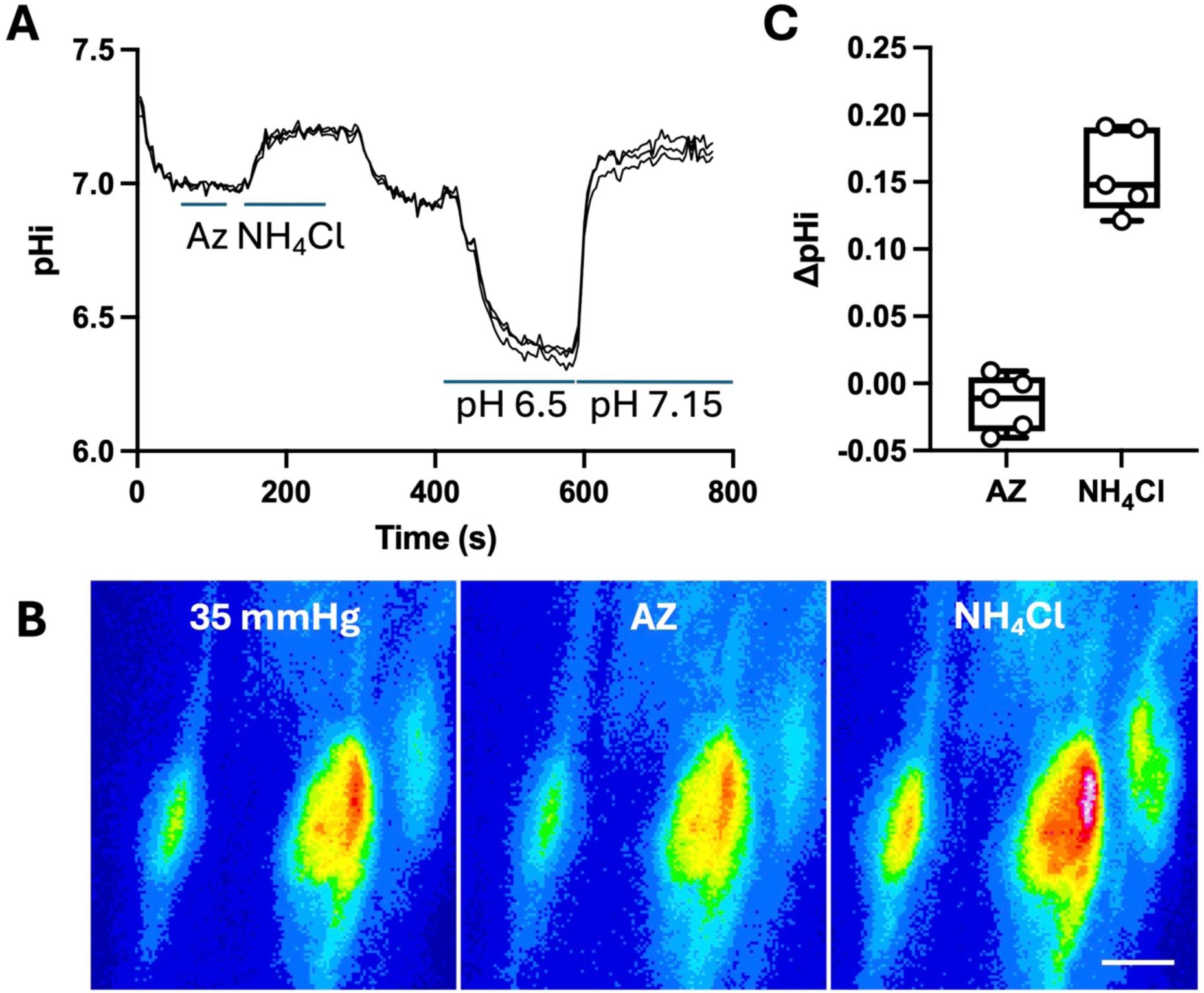
Effects of acetazolamide and NH_4_Cl on the intracellular pH of Schwann cell paranodes. A) representative trace showing the normalised BCECF fluorescence ΔF/F_0_ in response to changing pH to 6.5 and 7.15, 100μM acetazolamide (AZ) and 100 μM NH_4_Cl. B) representative images for the trace for the BCECF fluorescence in each condition. Scale bar = 5 µm. C) Summary graph of effect of AZ and NH_4_Cl on intracellular pH. Each point represents a nerve.

**Figure 9 figure supplement 1.**
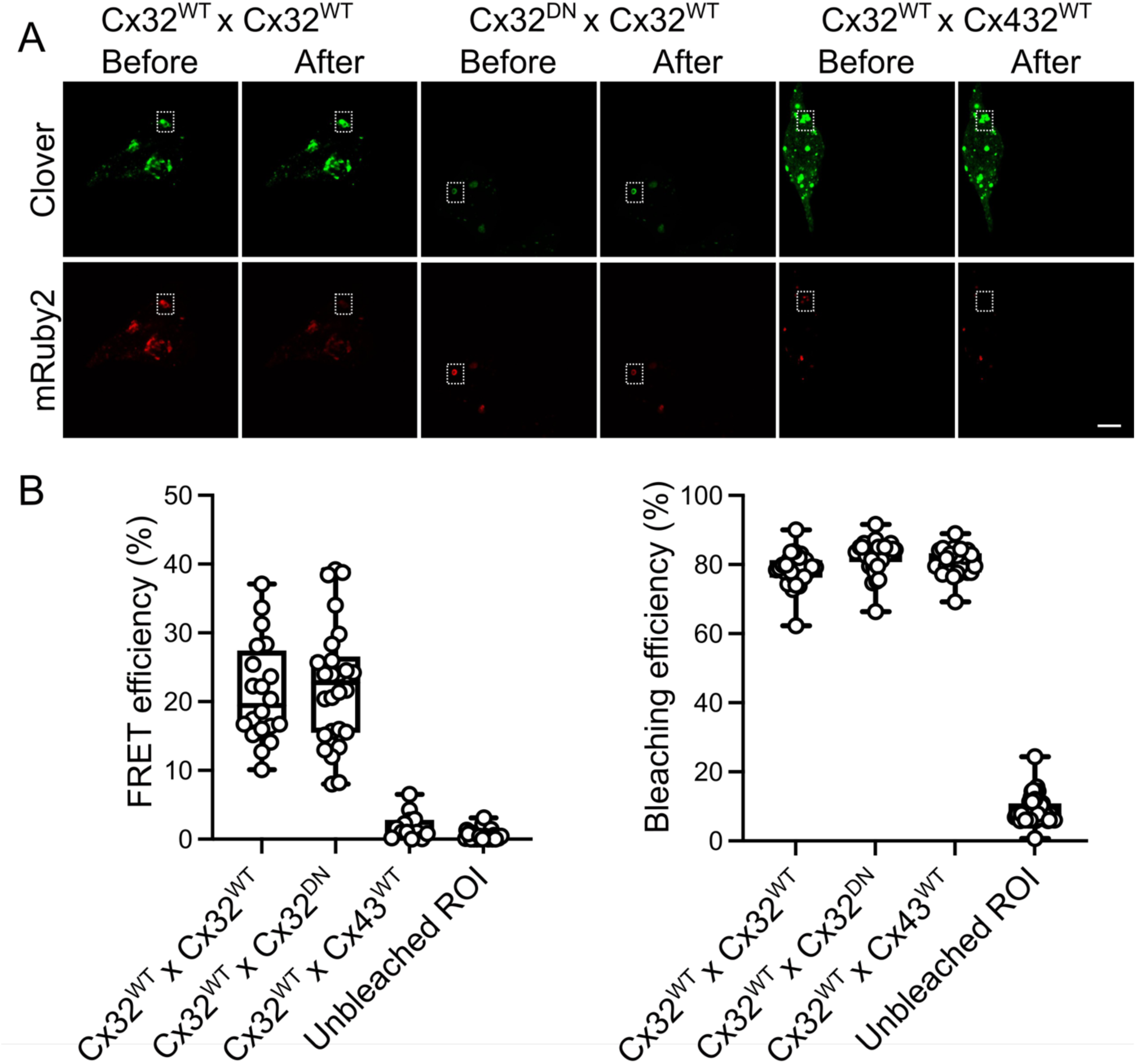
Cx32^DN^ coassembles with Cx32^WT^. A) Images of Clover and mRuby2 fluorescence before and after bleaching of the mRuby2 acceptor. The Clover fluorescence becomes brighter following bleaching for the Cx32^DN^ and Cx32^WT^ or Cx32^WT^ and Cx32^WT^ pairs but not for the Cx32^WT^ and Cx43^WT^ pair. B) Values for the FRET efficiency and a comparison of bleaching efficiency for the different pairings.

**Figure 9 figure supplement 2.**
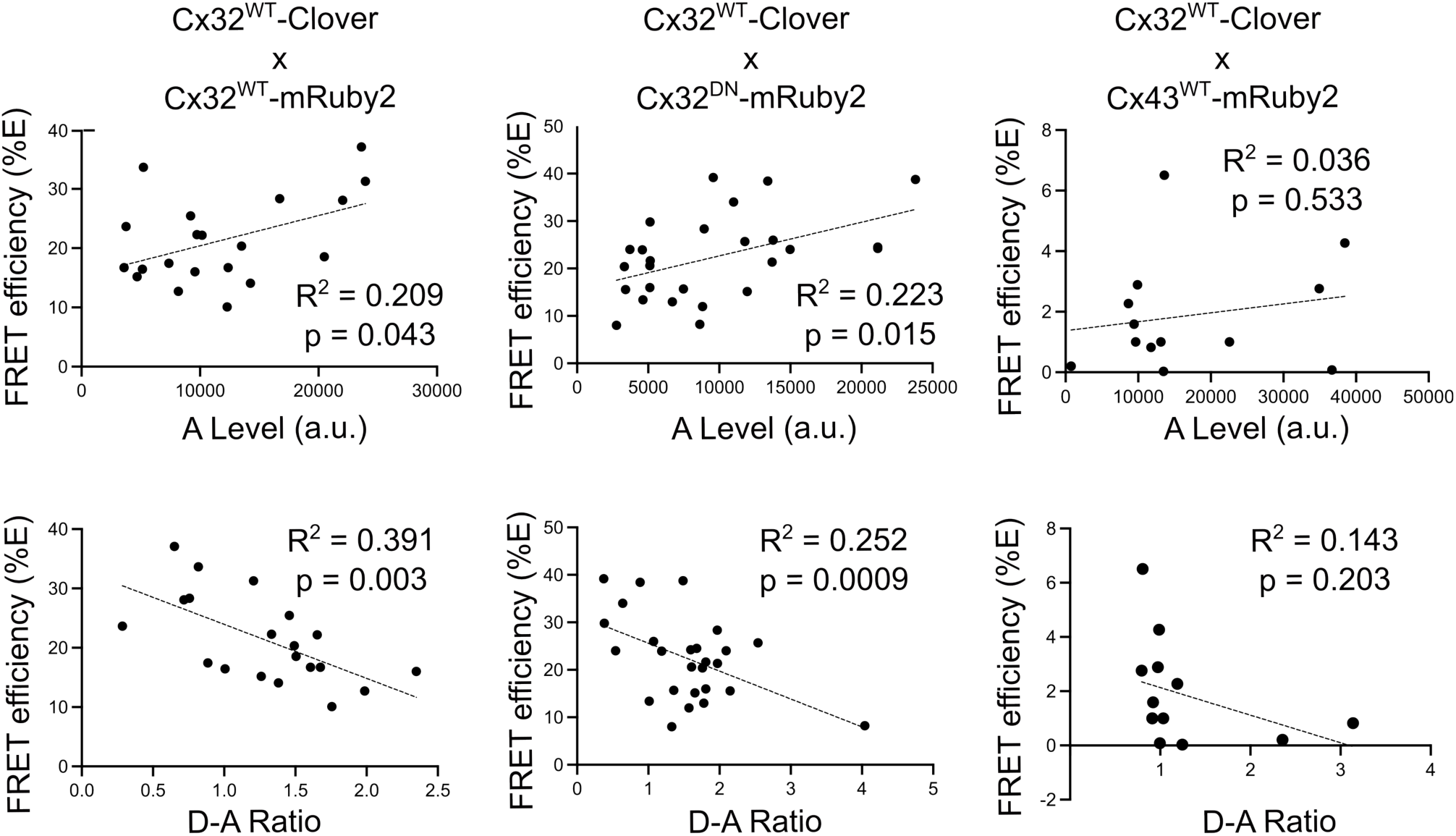
Cx32^DN^ coassembles with Cx32^WT^. The dependence of the FRET efficiency (E) on the Acceptor level (A) and Donor to Acceptor (D-A) Ratio. A negative correlation between E and D-A ratio indicates coassembly into hexamers as opposed to random association in the membrane.

**Figure 9 figure supplement 3.**
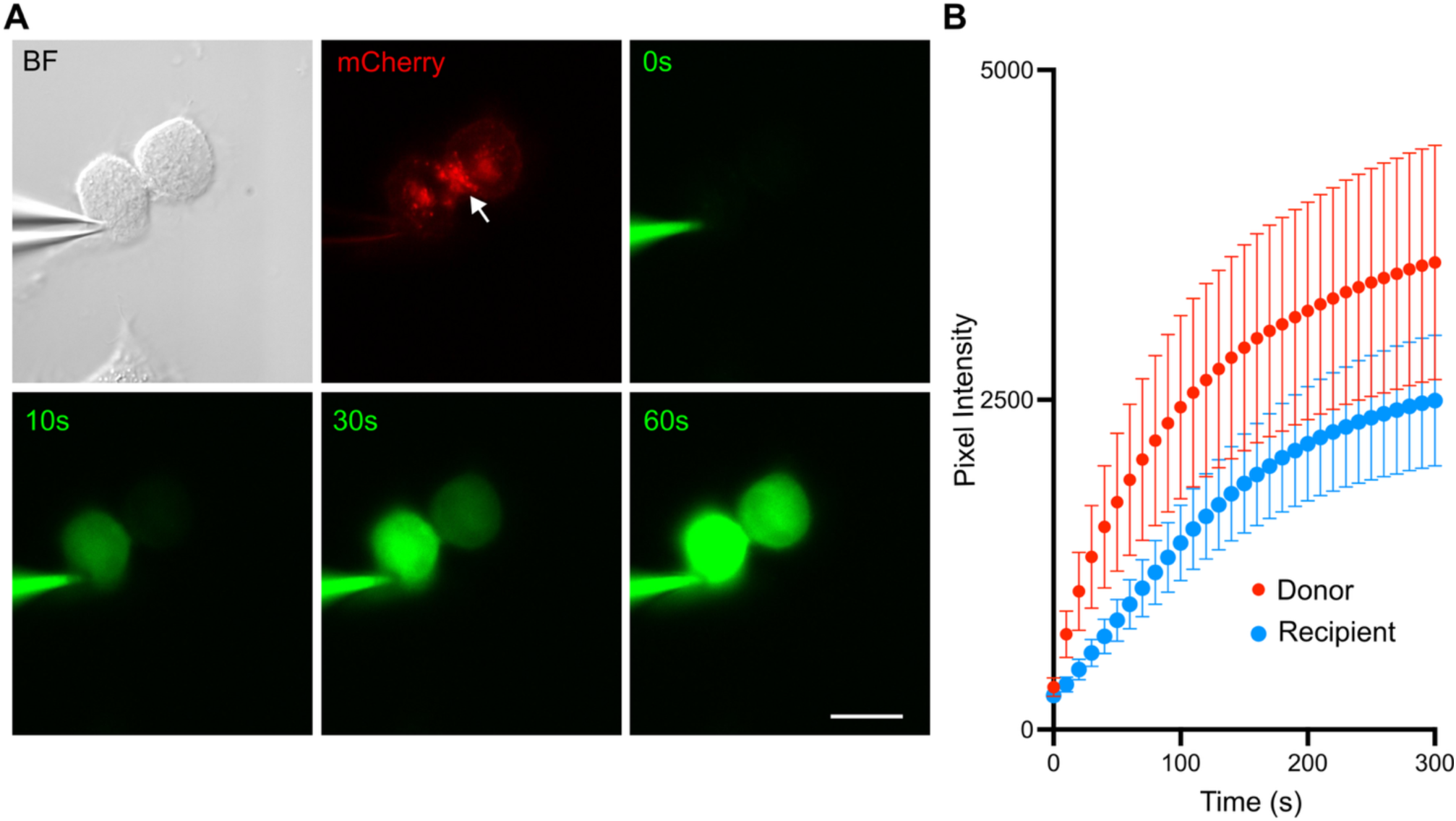
Cx32^DN^ forms functional gap junctions. A) Images of cells expressing Cx32 under brightfield (BF), the mCherry tag (red) and NBDG, a fluorescent glucose analogue (green). NBDG is present in the patch pipette and readily diffuses between the cells. Scale bar 20 µm. B) Changes in NBDG fluorescence over time in the Donor and Recipient cells showing the ready transfer of NBDG via Cx32^DN^ gap junction channels (n=8).

**Figure 10, figure supplement 1.**
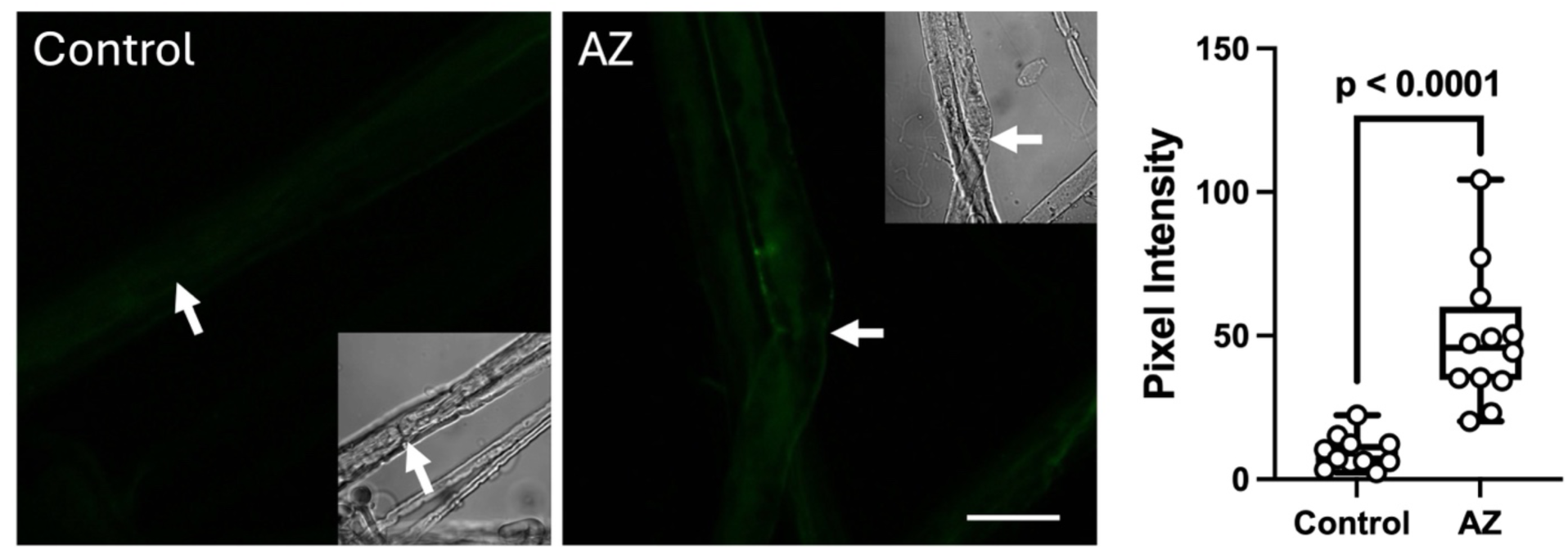
Inhibition of carbonic anyhydrase increases loading of FITC into paranodes and outer myelin in the absence of electrical stimulation. The images show the FITC fluorescence in the control and presence of 100 µM acetazolamide (AZ). The boxplot shows quantification of the fluorescence and that the increase in background fluorescence is significant (control vs AZ, MW test, p<0.0001). Each point represents a separate ROI from 5 different nerves.

**Figure 12 figure supplement 1.**
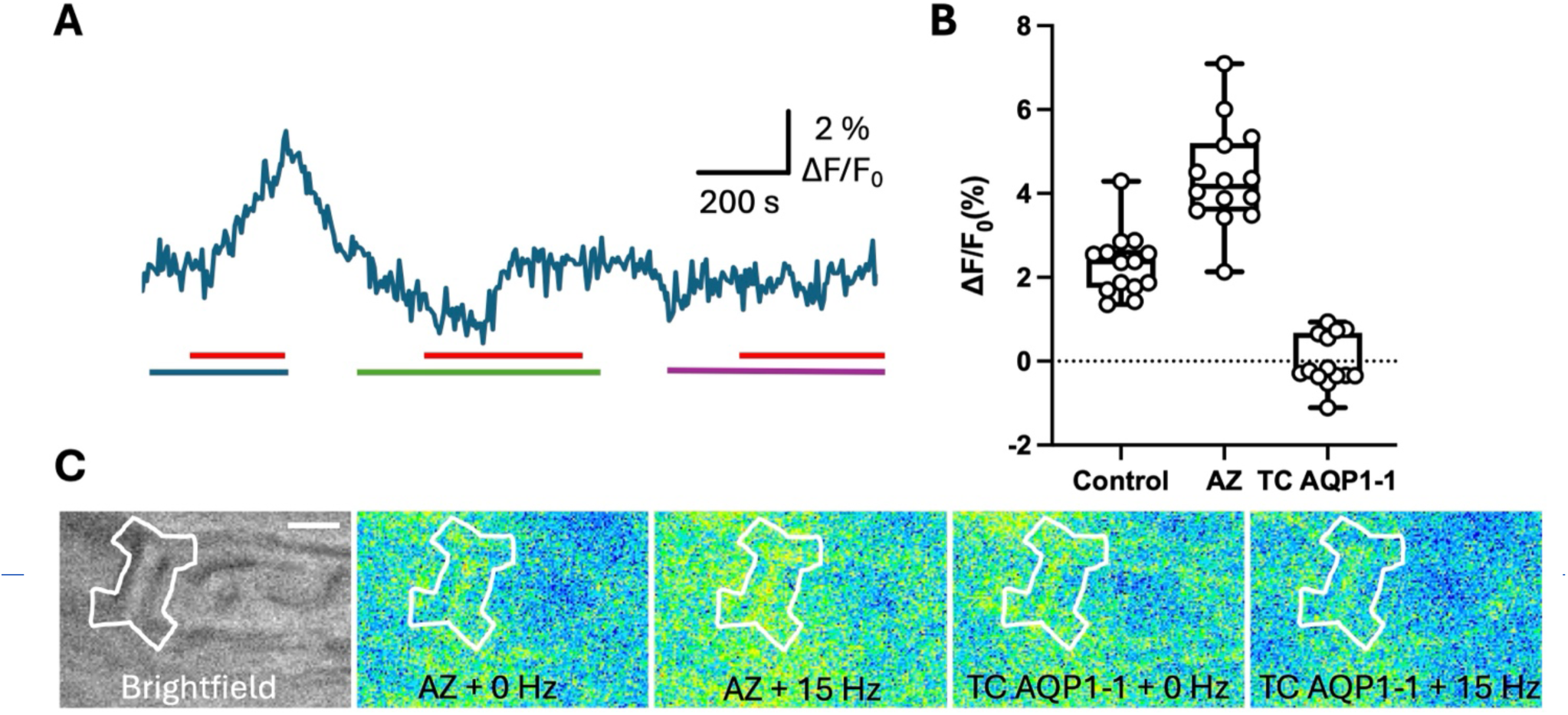
Activity dependent Ca^2+^ influxes into Schwann cell paranodes are dependent on CO_2_. A) Representative trace showing change in normalised Fluo4 fluorescence in response to 15 Hz electrical stimulation (red bar) in the presence of 100 µM acetazolamide (blue bar), in control aCSF (green bar), or in the presence of 80 µM TC AQP1-1 (purple bar). B) Boxplot showing the change in normalised fluorescence (ΔF/F_0_) evoked in Fluo4 loaded Schwann cell paranodes in response to high frequency electrical stimulation in each condition. Each datapoint consists of a paranode, with all the data collected from 4 sciatic nerves. Kruskal Wallis ANOVA, p < 0.0001. Pairwise MW comparisons: control vs AZ, p < 0.001; control vs TC AQP1-1, p<0.001. C) Brightfield and fluorescence images showing the activity dependent Ca^2+^ increase in a paranode in the presence of acetazolamide (AZ) and the lack of an activity dependent increase in Ca^2+^ in the presence of TC AQP1-1.

**Figure 13 figure supplement 1.**
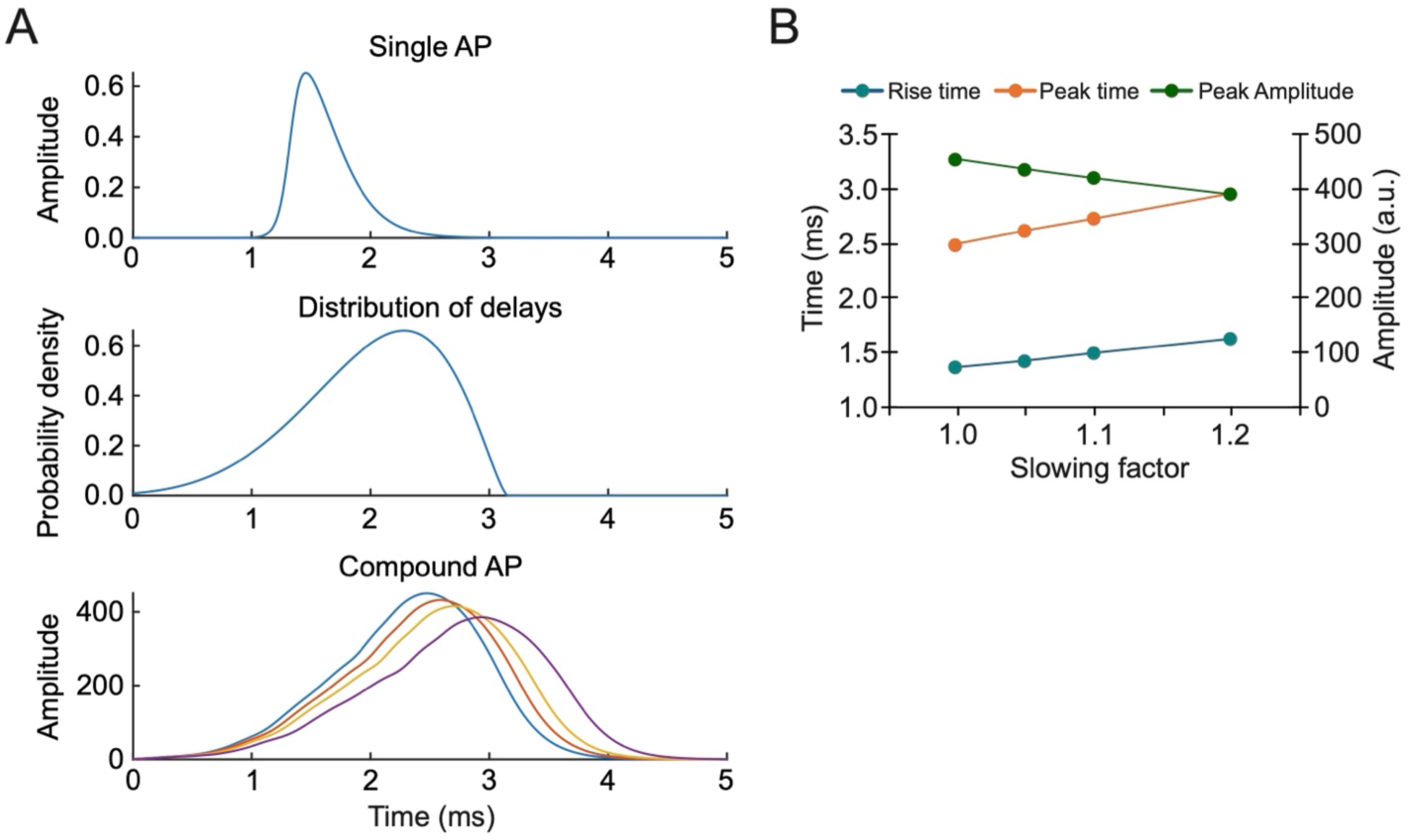
Model of the CAP to show how changing conduction velocity alters the rise time, time to peak and peak amplitude of the CAP. A) Shows the profile of an individual action potential (AP, top), the distribution of conduction delays (based on the distribution of diameters of fibres in the sciatic nerve (middle), and the CAP arising from summation of 2000 individual APs occurring with different conduction delays (bottom, light blue line) and three different slowing factors. B) Shows how the rise time, peak time and amplitude vary with the slowing factor. To mimic slowing, every conduction delay is multiplied by the slowing factor, thus when this is 1.0 no slowing occurs and when for example it is 1.2 each conduction delay is slowed by 20%. Matlab code for the model is provided in compoundAP.m.

