## Supplementary material for "Cell-to-cell signalling mediated via CO_2_: activity dependent axonal CO_2_ production opens Cx32 in the Schwann cell paranode": Matlab code for paranode model

%{

This specifies a model cell with a mitochondrion, a resting potential
(there is a K+ selective leak channel), and Cx32. CO2 produced by the
mitochondrion opens Cx32.

CO2 production is proportional to ATP production.

Carbonic anhydrase is present to convert CO2 to HCO3-.

Loading of a negatively charged dye (CBF) that can permeate Cx32 is 
calculated. FITC same as CBF.

%}


function dydt = CO2Cx32Model(t,y,Vm,Duration)

%Mito ATP production model based on Matsuda et al 2020 Quant biol 8, 288

%{

y(1) Y
y(2) Z
y(3) ATP
y(4) CO2
y(5) V
y(6) CBF  (FITC)

%}

Zbasal=8.5e-1;
Ybasal= 5e-2; % 9.9e-2; %5e-2;
Xlevel=0;
Kd1=0.77;
Kd2=1.8;
n1=0.95;
n2=1.5;
K1= 8e-2; % /min
K2= 12e-2; % 8.4e-3;  % /min

Tau1=1; %0.57;  %min
Tau2=4.3; % min

%CAII

KMca=13; %mM, from literature
VMca=Vm*60; %mM/min
Kspont=1.2e1; %spontaneous conversion 100 fold slower than CA


%stimulation

if (t>=30) && (t<=30+Duration) 
    Xlevel=30;


else
    Xlevel=0;
end


Yprod=Xlevel^n1/(Kd1+Xlevel^n1);
Yloss=(y(1)-Ybasal)/Tau1;
dY=Yprod-Yloss;

Zprod=Xlevel^n2/(Kd2+Xlevel^n2);
Zloss=(y(2)-Zbasal)/Tau2;
dZ=Zprod-Zloss;

ATPprod=K1*y(1);
ATPloss=K2*y(3)*y(2);
dATP=ATPprod-ATPloss;

%Calculating CO2 proportional to Y
CO2prod=2000*K1*y(1); % fudge to get mM levels
CO2loss=VMca*y(4)/(y(4)+KMca)+Kspont*y(4); %MM model
dCO2=CO2prod-CO2loss;

% Gating Cx32
HillCx=4;
Km=2.3;
Ecx=0;

CO2pow=(y(4)/Km)^HillCx;
gCx32=CO2pow/(1+CO2pow);
ICx32=20*gCx32*(y(5)-Ecx);


%K leak current

%{
% Nernstian version
gK=1;
Ek=-90; %mV
Ik=gK*(y(5)-Ek);
%}

%Current due to permeation of CBF (FITC)

Pcbf=1e-1;
CBFo=2e-1; %mM

%{
% Nernst with a valence of-1
%Ecbf=-59*log(CBFo/y(6));   
%Icbf=40*gCx32*Pcbf*(y(5)-Ecbf);
%%{
%}


% For GHK equation valence = -1
% This is more realistic than Nernst given unequal concentrations of CBF

F=96480;
R=8.314;
T=298;
FRT=F/(R*T);

V=y(5)/1000;  % convert to V for GHK

gK=1.5E-5; %GHK version of leak, probably more realistic
if V~= 0  %this is to avoid division by zero when V=0
    Ik=gK*V*F*FRT*(120-3*exp(-V*FRT))/(1-exp(-V*FRT));%GHK version of leak [K] in mM
    Icbf=gCx32*Pcbf*V*F*FRT*(y(6)-CBFo*exp(V*FRT))/(1-exp(V*FRT));
else
    Ik=gK*F*117; %(120-3)
    Icbf=gCx32*Pcbf*F*(y(6)-CBFo);
end


Jcbf=Icbf/F;

dCBF=Jcbf;

dV=(-Ik-ICx32-Icbf);

dydt = [dY;dZ;dATP;dCO2;dV;dCBF];

end
